# β-defensin gene copy number variation in cattle

**DOI:** 10.1101/2024.06.24.600336

**Authors:** Ozge Sidekli, John Oketch, Sean Fair, Kieran G. Meade, Edward J. Hollox

## Abstract

β-defensins are peptides with antimicrobial roles, characterized by a conserved tertiary structure. Beyond antimicrobial functions, they exhibit diverse roles in both the immune response and fertility, including involvement in sperm maturation and function. Copy number variation (CNV) of β-defensin genes is extensive across mammals, including cattle, with possible implications for reproductive traits and disease resistance. In this study, we comprehensively catalogue 55 β-defensin genes in cattle. By constructing a phylogenetic tree to identify human orthologues and lineage-specific expansions, we identify 1:1 human orthologues for 35 bovine β-defensins. We also discover extensive β-defensin gene CNV across breeds, with *DEFB103* in particular showing extensive multiallelic CNV. By comparing β-defensin expression levels in testis from calves and adult bulls, we find that 14 β-defensins, including *DEFB103*, increase in expression during sexual maturation. Analysis of β-defensin gene expression levels in the caput of adult bull epididymis, and β-defensin gene copy number, in 94 matched samples shows expression level of four β-defensins are correlated with genomic copy number, including *DEFB103*. We therefore demonstrate extensive copy number variation in bovine β-defensin genes, in particular *DEFB103*, with potential functional consequences for fertility.

## Introduction

One of the largest sub-classes of the defensin protein family, β-defensins were first characterised as short cationic peptides produced by many multicellular organisms with an antimicrobial role (Hollox and Abujaber, 2017; Shelley et al., 2020). β-defensins peptide sequences are very diverse, with conservation of a core glycine and aspartic acid residue, and six cysteines which form three disulphide bridges in the mature protein leading to a conserved tertiary structure characteristic of all β-defensins (Cormican et al., 2008). Secreted at epithelial and mucosal surfaces, it has become increasingly clear that defensins in fact have adopted a multitude of additional roles over the course of their evolution. Evidence now shows that defensins interact with other components of the innate and adaptive immune system, in often complex and contradictory ways (Shelley et al., 2020). For example, human β-defensin 3 (hBD-3) has dual roles in modulating the immune response. It can directly act as a chemoattractant for monocytes by binding to the CCR2 receptor, similar to traditional chemokines (Rohrl et al., 2010; Wu et al., 2003). Additionally, hBD-3 can indirectly promote a proinflammatory response by stimulating the production of various cytokines and chemokines from other cells, thereby amplifying the recruitment and activation of immune cells (Niyonsaba et al., 2010).

β-defensins have roles beyond modulation of the immune response. It is clear, that, in humans at least, some β-defensins are expressed in different regions of the epididymis post-puberty, with many only being expressed in the epididymis (Rodriguez-Martinez, 2003; Yamaguchi et al., 2002). Empirical evidence has implicated specific β-defensins in sperm maturation, function and ultimately regulating male fertility (Dorin, 2015; Zhou et al., 2013). The β-defensin best characterised in its role in male fertility is β-defensin 126 (*DEFB126*) which has been extensively characterised in humans and macaques (Aram et al., 2020). *DEFB126* protein is secreted by the epididymis, is highly glycosylated, and subsequently adsorbed onto the surface of sperm as they travel through the epididymis. Through the addition of a negative charge, *DEFB126* facilitates penetration of cervical mucus by the sperm and possibly protects the sperm from female immune recognition (Tollner et al., 2008). The adsorbed *DEFB126* is subsequently shed during capacitation to allow fertilisation to occur (Tollner et al., 2012, 2011). DEFB126 has also been shown to affect sperm motility in cattle via the enhancement of sperm binding to oviductal epithelium (Lyons et al., 2018) and the prevention of sperm agglutination (Fernandez-Fuertes et al., 2016). Other β-defensins have been shown to affect fertility, and given their widespread expression in the male reproductive tract, much remains to be discovered about the functional role of these proteins (Dorin and Barratt, 2014; Solanki et al., 2023). Beyond *DEFB126*, most knowledge so far is from analysis of β-defensins and fertility in rodents. A mouse knockout of a cluster of nine β-defensins in mice renders male mice infertile (Zhou et al., 2013), with evidence showing sperm defects to centrioles etc.

Across mammals, different members of the β-defensin gene family show extensive copy number variation (CNV) between individuals (Hollox and Abujaber, 2017). This has been best-characterised in humans (Hollox et al., 2003; Logsdon et al., 2021), but CNV has been characterised in detail in macaque (Ottolini et al., 2014) and has been detected in genome-wide scans in dogs (Leonard et al., 2012), pigs (Wang et al., 2013), chickens (Lee et al., 2016), and cattle (Bickhart et al., 2012). CNV of the β-defensin gene *DEFB4* has phenotypic consequences in humans, as it has been shown to be associated with its encoded protein levels and mucosal antimicrobial activity in the cervix of the CNV (James et al., 2018), and associated with the risk of the inflammatory skin disease psoriasis (Stuart et al., 2012). However, a robust association of β-defensin CNV with a phenotype outside humans has not yet been shown, nor has an association been shown with a phenotype reflecting the reproductive role of β-defensins rather than their antimicrobial or inflammatory roles.

In this study, we have focused on characterising the CNV of β-defensins in domestic cattle (Bos *taurus*). For the dairy industry, in particular, a focus on improving food security though higher milk yields has led to a significant decline in fertility (Makanjuola et al., 2020). Fertility is a critical trait for sustainable animal production systems and significant variation in bull fertility can account for low pregnancy rates of individual bulls, and also, although multiple SNPs have been associated with fertility, the ability to predict poor bull fertility remains elusive despite the extensive use of sperm motility and morphology quality control checks prior to the release of semen into the field (Berry et al., 2014; Fair and Lonergan, 2018). Together with the emergence of artificial insemination (AI) activities, this has renewed the interest and importance of understanding the role of genetic variation in reproductive traits, including bull subfertility. Therefore, understanding the extensive genetic variation of the β-defensin family, extremely important in mammalian reproduction, in cattle, is an important step in understanding its phenotypic consequences for reproduction.

In cattle, 57 β-defensins have been previously identified based on sequence similarity, including the genes encoding tracheal antimicrobial peptide (*TAP*) and lingual antimicrobial peptide (*LAP*) (Cormican et al., 2008). These β-defensin genes occur in four clusters on chromosome 8 (cluster A), chromosome 13 (cluster B), chromosome 23 (cluster C) and chromosome 27 (cluster D), orthologous to the human β-defensin clusters on chromosome 8 (proximal), chromosome 20, chromosome 6 and chromosome 8 (distal) (Patil et al., 2005). By mapping whole genome short-read sequencing to a cattle genome assembly, eleven β-defensins were reported to show copy number variation, with average copy numbers suggesting extensive gene duplication (Bickhart et al., 2012).

The development of cattle genomic resources, and improvement of the cattle reference genome, has important consequences for the accuracy of CNV inference. Bickhart et al (2012) used Btau_4.0 as the reference genome, and the subsequent genome assembly (UMD3.1) allowed incorporation of previously-unplaced genomic contigs. The most recent assembly (ARS-UCD1.2), which incorporates long sequencing reads from Pacific Biosystems technology, represents a 200-fold improvement in sequence contiguity and a 10-fold improvement in sequence accuracy per-base, compared to UMD3.1 (Rosen et al., 2020). This improved contiguity will have particularly improved regions with segmental duplications prone to CNV, which are known to be challenging to assembly using just short-read sequences. Whole-genome short-read sequencing data provided by the 1000 Bulls consortium has allowed an assessment of the genome-wide sequence variation between and within cattle breeds (Hayes and Daetwyler, 2019). Short-read sequencing data can also be used to determine the nature and extent of CNV in a genome. Taken together, these genomic resources provide an ideal starting point for a focused analysis on the CNV of β-defensins in cattle.

In this work, comprehensively catalogue the β-defensin genes in the latest high-contiguity bovine genome assembly, and by constructing a phylogeny identify human orthologues and recent β-defensin expansions in the bovine lineage.

We identify the range and extent of CNV in β-defensin genes across 7 different (Holstein, Hereford, Charolais, Limousin, Simmental, Angus, Swiss Braunvieh and their cross breeds) breeds, focusing particularly on Holsteins, the primary dairy breed worldwide. We develop digital droplet PCR (ddPCR) assays to extend this analysis to unsequenced bulls and measure copy numbers of Holstein and Jersey bulls in the Irish Dairy herd. We then analyse the expression levels of β-defensin genes in the caput epididymis and the relationship of CNV with gene expression.

## Methods

### 1000 Bulls samples

Whole genome sequences (WGS) were downloaded from 1000 Bulls Consortium data at the European Nucleotide Archive (ENA, https://www.ebi.ac.uk/ena), focusing on Holstein-Friesian (n=30), but also including Hereford (n=6), Charolais (n=26), Limousin (n=19), Simmental (n=9), and Angus (n=10) breeds, (Purfield et al., 2019), (Supplementary Table 1).

### Other data sources

RNA-seq data for Holstein (adult, n = 24) and Hereford (calf, n=3; adult, n =7) were obtained from ENA under accession numbers PRJNA566324, PRJNA760322, PRJNA263600, PRJNA379574 and PRJNA526257 (Supplementary Table 2). Data from the Swiss Braunvieh and their cross breeds (n=103), used for correlation analysis of CNV with gene expression at transcript level in caput epididymis, were generated by Mapel et al (2024) (Supplementary table 3). From this dataset, a total of 9 samples were removed from the study, seven due to insufficient coverage in the DNA sequence and two due to corrupt raw files or missing second reads in the RNA sequence. In total, 94 samples were retained for analysis, encompassing both transcript-level gene expression and CNV assessment. These datasets are accessible on the ENA under the accession number PRJEB46995 (Supplementary Table 3).

### DNA samples and genomic DNA extraction

Genomic DNA was extracted using a standard phenol-chloroform method from semen from Irish Holstein-Friesian bulls (n=20) and Jersey bulls (n=15) and positive control bulls (n=4) selected from WGS analysis. The genomic DNA concentration was determined by measuring the fluorescence signal using the Qubit 2.0 Fluorometer.

### β-defensin search and alignment

Using keywords "Human", "Bovine", and "β-defensin; DEFB", β-defensin gene and β-defensin predicted protein sequences were retrieved from Ensembl (Hubbard et al., 2002), National Center for Biotechnology Information (NCBI; (Pruitt et al., 2014)), UniProt (https://www.uniprot.org/; (Uniprot Consortium, 2021)) and University of California Santa Cruz (UCSC) Genome Browser (https://genome.ucsc.edu/; (Kent et al., 2002)) by querying the reference genomes ARS-UCD1.2 for cattle and GRCh38 for human.

A total of 66 bovine and 39 human β-defensin gene sequences and predicted proteins sequences were identified. To refine the dataset, the signal peptides of the protein sequences were cleaved using resources from the UniProt website, and identical sequences, those that did not map to the latest reference genome, and those lacking a predicted start codon were excluded from the analysis. A curated dataset consisting of 55 β-defensin predicted protein sequences in cattle (Supplementary table 4) and 37 β-defensin predicted protein sequences in humans (Table 1) were aligned using Clustal Omega (Madeira et al., 2022) .

**Table 1.** Human orthologs of bovine β-defensin genes.

| Ensembl Gene ID | Symbol | Location ARS-UCD1.2 assembly | Protein | Protein Accession | Orthologue |
| --- | --- | --- | --- | --- | --- |
| Not Available | *DEFB43* | chr8:7,449,722-7,456,717 | BBD43 | XP_024852154.1 | HBD131A |
| ENSBTAG00000005<br>4255 | *DEFB134* | chr8:7,469,680-7,473,928 | BBD134 | XP_024852155.1 | HBD134 |
| ENSBTAG00000005<br>1632 | *DEFB136* | chr8:7,499,869-7,500,896 | BBD136 | XP_024852156.1 | HBD136 |
| ENSBTAG00000004<br>9760 | *DEFB132-*<br>*1* | chr13:60,755,471-60,757,652 | BBD132<br>-1 | XP_005215029.4 | HBD132 |
| ENSBTAG00000004<br>8288.2 | *DEFB129-*<br>*2* | chr13:60,772,966-60,775,142 | BBD129<br>-2 | ENSBTAT000000064<br>705.2 | HBD129 |
| ENSBTAG00000005<br>1173 | *DEFB128* | chr13:60,785,898-60,787,480 | BBD128 | ENSBTAT000000068<br>291.1 | HBD128 |
| ENSBTAG00000005<br>4958 | *DEFB127* | chr13:60,793,742-60,796,520 | BBD127 | XP_005215144.1 | HBD127 |
| ENSBTAG00000005<br>1033 | *DEFB126* | chr13:60,806,515-60,811,042 | BBD126 | ENSBTAT000000071<br>785.1 | HBD126 |
| Not Available | *DEFB125*<br>*A* | chr13:60,828,555-60,835,043 | BBD125<br>A | XP_024856951.1 | HBD125 |
| ENSBTAG00000005<br>3093 | *DEFB125* | chr13:60,828,987-60,829,208 | BBD125 | G8CY10 | HBD125 |
| ENSBTAG00000004<br>9491 | *DEFB115* | chr13:60,873,647-60,884,399 | BBD115 | XP_024857141.1 | HBD115 |
| ENSBTAG00000005<br>0556 | *DEFB29* | chr13:60,893,940-60,905,009 | BBD29 | XP_002692327.1 | None |
| ENSBTAG0000005<br>2478 | *DEFB116* | chr13:60,920,317-60,925,563 | BBD116 | XP_010809939.1 | HBD116 |
| ENSBTAG0000004<br>8009 | *DEFB117* | chr13:60,956,394-60,959,265 | BBD117 | ENSBTAT00000063<br>334.2 | HBD118 |
| ENSBTAG0000005<br>2743 | *DEFB118* | chr13:60,979,467-60,979,700 | BBD118 | G8CY17 | HBD118 |
| Not Available | *DEFB119-*<br>*1* | chr13:60,981,163-60,990,988 | BBD119<br>-1 | NP_001095131.1 | HBD119 |
| ENSBTAG0000000<br>3364 | *DEFB119-*<br>*2* | chr13:60,989,366-60,990,988 | BBD119<br>-2 | NP_001095829 | None |
| ENSBTAG0000004<br>8949 | *DEFB121* | chr13:61,007,870-61,009,276 | BBD121 | XP_005215031.1 | HBD121 |
| ENSBTAG0000002<br>7384 | *DEFB122*<br>*A* | chr13:61,019,572-61,023,625 | BBD122<br>A | NP_001095809.1 | None |
| ENSBTAG0000002<br>7383 | *DEFB122* | chr13:61,030,365-61,034,955 | BBD122 | NP_001071575.1 | None |
| Not Available | *DEFB123-*<br>*2* | chr13:61,038,268-61,052,092 | BBD123<br>-2 | XP_024856654.1 | HBD123 |
| ENSBTAG0000002<br>0555 | *DEFB123-*<br>*1* | chr13:61,041,375-61,051,806 | BBD123<br>-1 | NP_001095813.1 | HBD123 |
| ENSBTAG0000003<br>1254 | *DEFB124* | chr13:61,067,234-61,069,998 | BBD124 | NP_001095830.1 | HBD124 |
| ENSBTAG0000005<br>4219 | *DEFB114* | chr23:22,612,721-22,615,376 | BBD114 | XP_024839622.1 | HBD114 |
| ENSBTAG0000005<br>4396 | *DEFB113* | chr23:22,632,551-22,635,067 | BBD113 | XP_024839712.1 | HBD113 |
| Not Available | *DEFB110-*<br>*1* | chr23:22,644,099-22,656,635 | BBD110<br>-1 | XP_024839746.1 | HBD110 |
| ENSBTAG0000005<br>1453 | *DEFB110-2* | chr23:22,651,406-22,656,635 | BBD110<br>-2 | XP_002697354.1 | None |
| ENSBTAG0000004<br>6711 | *DEFB112* | chr23:22,662,983-22,671,406 | BBD112 | XP_024839747.1 | HBD112 |
| ENSBTAG0000005<br>0611 | *DEFB1-2* | chr27:5,960,716-5,968,525 | BBD1-2 | ENSBTAT00000068<br>469.1 | None |
| ENSBTAG0000004<br>8171 | *TAP* | chr27:6,013,830-6,015,648 | TAP | NP_777201.1 | None |
| ENSBTAG0000005<br>2579 | *DEFB103B* | chr27:6,023,993-6,025,303 | BBD103<br>B | XP_010818521.1 | HBD103B/HBD-Like |
| ENSBTAG0000003<br>4952 | *SPAG11B* | chr27:6,045,534-6,050,941 | BSPAG1<br>1B | XP_005225990.1 | H-SPAG11B |
| ENSBTAG0000004<br>8847 | *DEFB104* | chr27:6,069,928-6,074,858 | BBD104 | ENSBTAT00000075<br>330.1 | HBD104A |
| ENSBTAG0000004<br>9070 | *DEFB106A* | chr27:6,076,779-6,079,838 | BBD106 | ENSBTAT00000075<br>644.1 | HBD106B-2 |
| Not Available | *DEFB15* | chr27:6,076,989-6,079,896 | BBD15 | XP_024842096.1 | HBD106B-2 |
| ENSBTAG0000005<br>2911 | *DEFB105* | chr27:6,082,209-6,084,799 | BBD105 | XP_015316308.1 | HBD105A |
| ENSBTAG0000005<br>4607 | *DEFB107A* | chr27:6,090,910-6,095,691 | BBD107<br>A | XP_002698669.1 | HBD107A/HBD1<br>06B-1 |
| ENSBTAG0000005<br>3557 | *DEFB4A* | chr27:7,138,963-7,140,882 | BBD4A | NP_777200 | None |
| Not Available | *DEFB5* | chr27:7,139,081-7,140,848 | BBD5 | NM_001130761.1 | None |
| ENSBTAG0000003<br>3545 | *EBD* | chr27:7,165,176-7,180,422 | EBD | NP_783634 | None |
| ENSBTAG0000005<br>1223 | *DEFB130-1* | chr27:6,192,946-6,196,744 | BBD130 | XP_024842219.1 | HBD130A |
| ENSBTAG0000005<br>3889 | *LAP* | chr27:6,234,006-6,235,870 | LAP | NP_982259.3 | None |
| ENSBTAG0000004<br>6937 | *DEFB33* | chr27:6,238,997-6,242,941 | BBD33 | NP_001161717 | None |
| ENSBTAG0000005<br>0630 | *DEFB13* | chr27:6,326,696-6,328,616 | BBD13 | NP_001311470.1 | None |
| ENSBTAG0000005<br>0419 | *DEFB103A* | chr27:6,432,334-6,433,571 | BBD103<br>A | XP_015316312.1 | HBD103B/HBD-Like |
| ENSBTAG0000004<br>8737 | *DEFB10* | chr27:6,596,623-6,598,299 | BBD10 | NP_001108556.1 | None |
| Not Available | *DEFB3* | chr27:6,676,309-6,723,692 | BBD3 | NP_001269510 | None |
| ENSBTAG0000005<br>1383 | *DEFB7* | chr27:6,676,383-6,678,273 | BBD7 | NP_001095832.1 | None |
| Not Available | *DEFB6* | chr27:6,676,387-6,996,057 | BBD6 | NP_001300920 | None |
| Not Available | *DEFB9* | chr27:6,676,540-6,678,161 | BBD9 | P46167 | None |
| ENSBTAG0000005<br>2312 | *DEFB1-1* | chr27:6,721,772-6,723,692 | BBD1-1 | NP_001311473.1 | HBD1 |
| ENSBTAG0000005<br>3456 | *DEFB402* | chr27:6,855,327-6,856,946 | BBD402 | Q5W5I5 | None |
| ENSBTAG0000005<br>4169 | *DEFB108B* | chr27:7,272,455-7,275,948 | BBD108<br>B | XP_024842173.1 | HBD108B |
| Not Available | *DEFB109* | chr27:7,281,838-7,289,252 | BBD109 | XP_024842216.1 | HBD109A |

### Phylogenetic analysis

The multiple protein sequence alignment of bovine (n=54; BBD129-1 was omitted due to the same mature protein sequence as BBD129-2) and human β-defensins (n=37) was used to construct a phylogenetic tree by maximum-likelihood using IQ-TREE v1.6.12 (Nguyen et al., 2015), using the JTTDCMut+I+G4 model (revised JTT matrix, allowing for invariable sites and a discrete 4-state gamma model for site rate heterogeneity), and 1000 bootstrap replicates.

### Copy number analysis using whole genome short read sequences

Fastq reads were aligned to the reference genome ARS-UCD1.2 using BWA-mem2 v 2.2.1 (Vasimuddin et al., 2019). Alignment quality was assessed using QualiMap, v 2.3 (Okonechnikov et al., 2016). Mapped genomes were sorted using SAMtools (v1.17) to generate BAM files. PCR duplicates were removed using Picard v 2.6.0 (https://broadinstitute.github.io/picard/). Three samples (SRR1262661, SRR1262663 and SRR1262668) were excluded as their mean coverage fell below a threshold of 10x.

Genomic locations of the 55 β-defensin genes analysed were converted into a .bed file, and a custom Perl script counted mapped sequence reads for the regions defined in the .bed file. Read counts were normalized against a reference gene (*TP53*), then normalized by dividing by the average number of reads for each gene, then multiplied by two to get an estimate of copy number per diploid genome. Threshold values of <1.5 and >2.5 were used to identify deletions and duplications, respectively based on the expected copy number of 2 for the normal diploid genome. Three copies of the *DEFB103* gene are assembled in ARS-UCD1.2 (*DEFB103A*, *DEFB103A*-like and *DEFB103B*), so an expected copy number of 6 was used for the normal diploid genome.

### Copy number analysis using digital droplet PCR (ddPCR)

A subset of 31 β-defensin genes with previous evidence suggesting a link with fertility were selected (Supplementary table 5), including the multicopy *DEFB103*, with a single primer pair designed to bind to all three annotated *DEFB103* genes. PCR primers were designed to amplify products between 160-280 base pairs for 31 selected β-defensin genes and 80 base pairs for the reference gene *TP53*. Primers were validated using the in silico PCR tool in the UCSC Genome Browser to ensure that they were unique to the target sequence and that common sequence variants were absent (Supplementary table 5).

ddPCR was performed using the EvaGreen Supermix QX200™ Droplet Digital PCR (Bio-Rad Laboratories) according to manufacturer’s instructions, with QuantaSoftTM v1.7 and R v4.1.0, used to calculate the normalized copy number. In this assay, excluding *DEFB103*, a normal diploid CN of 2 was used as a reference, and thresholds of <1.5 and >2.5 were used to identify deletions and duplications, respectively. A diploid copy number of 6 was used as the reference for *DEFB103*, and distinct threshold values of <4.5 and >7.5 were employed to identify low copy number and high copy number, respectively, of this gene.

## RESULTS

### Genomic organisation of β-defensin genes in cattle

Our first aim was to infer the evolution of bovine β-defensin genes and identify the closest human orthologue, if possible. We identified 55 bovine predicted annotated β-defensin proteins and 37 predicted annotated human β-defensin proteins. Genes encoding the bovine proteins were annotated to one position on the ARS-UCD1.2 genome assembly, except BBD103, which was annotated three times to three separate regions. Aligning the predicted proteins (BBD103A, BBD103A-like, and BBD103B) shows that BBD103B is most similar to human β-defensin 103, with BBD103A encoding an extra 14-amino acid region at the N-terminus, likely to disrupt effective processing and secretion of any protein, and, and BBD103A-like lacking an initial methionine. For phylogenetic analysis, we used the two predicted full-length proteins.

An unrooted maximum-likelihood phylogenetic tree from predicted protein sequences shows that the bovine β-defensins cluster in the tree broadly according to their genomic clusters, indicating a pattern of gene duplication and divergence at those clusters (Figure 1). There are three exceptions – BBD134, BBD104 and BBD136, whose positions on the phylogenetic tree are incongruous compared to their cluster position of their encoding genes in the genome. The bootstrap values supporting their position in the tree are high, suggesting that this is not an error in the tree, and given the high quality of the reference genome, this points to an ancestral rearrangement after duplication and divergence of the β-defensin clusters.

**Figure 1.**
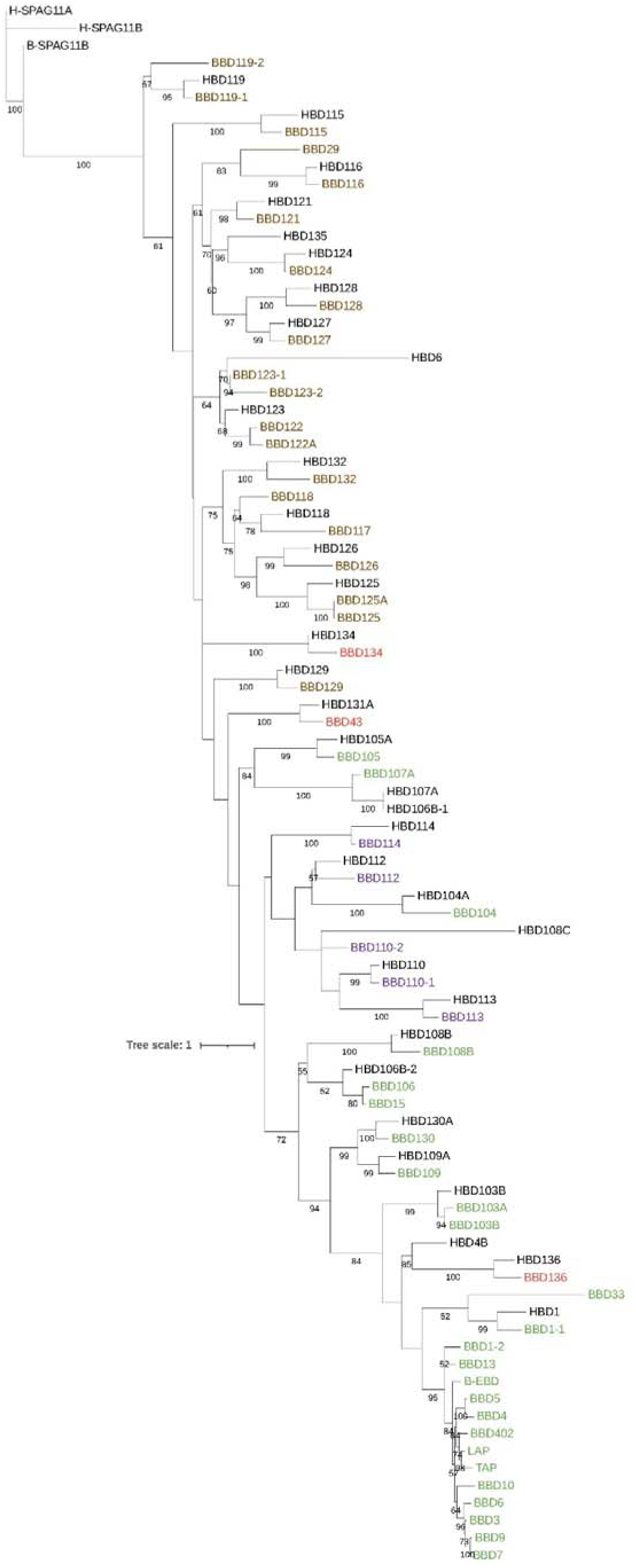
Phylogenetic tree of human and bovine β-defensin proteins. Bovine protein names are coloured according to the chromosomal location of their genes (brown: chromosome 13, green: chromosome 27, red: chromosome 8, purple: chromosome 23). Tree scale is indicated.

The phylogenetic tree also shows that 35 bovine β-defensins have a human orthologue (Table 1), with a cluster of 13 genes on chromosome 27 (cluster D), including the tracheal antimicrobial peptide and lingual antimicrobial peptide having no one-to-one human orthologues and being closely related to each other, suggesting duplication and divergence of this group after the divergence of primates and artiodactyls. The orthologues of the three bovine β-defensins that show incongruity between tree position and genomic cluster position of their gene (BBD134, BBD104 and BBD136) have human orthologues whose genes are also incongruent with their genomic cluster, suggesting that any evolutionary rearrangement occurred prior to primate-artiodactyl divergence.

### Analysis of copy number variation of β-defensins in cattle

To understand the CNV of β-defensin genes in cattle, we used a gene-focused approach to determine gene copy number from short-read sequencing alignments for 191 cattle across seven breeds. We found evidence of CNV for all the genes analysed (Supplementary figure 1), with diploid copy number usually ranging from 1 (heterozygote deletion) to three (heterozygote duplication) (supplementary table 2). There are exceptions – for example *DEFB103* is extensively multiallelic and shows a high copy number range of 1 to 29 (Supplementary figure 2), and *DEFB402* ranges from complete absence to 5 copies. This emphasises that, in most breeds, deletions and duplications of the genes are relatively uncommon, such that homozygous individuals are unusual. For most genes, some breeds did not show any copy number change – for example in Angus 34 β-defensins did not show CNV, although this is in a relatively small sample set of 10 bulls.

In contrast, the Holstein breed (n=27) and Swiss Braunvieh breed and their crosses (n=94) showed CNV of all β-defensins, suggesting that sample size of other breeds has limited the ability to detect rarer copy number variants.

In order to investigate whether any genes were on a copy number variable region together, we tested the degree to which CNV across all pairs of genes covaried in the WGS dataset. The rationale for this is that if two genes are on the same copy number variable region, then, across the 191 cattle, loss or gain of sequence read depth will be strongly correlated. By calculating the correlation coefficient between every pair of genes, and using a threshold of 0.9, we found three pairs of genes to covary : *DEFB7* and *DEFB9* (r=1.00), *DEFB15* and *DEFB106A* (r=0.99), and, unsurprisingly because they are alternative transcripts from the same genomic region, *DEFB110.1* and *DEFB110.2* (r=0.97). *DEFB7* and *DEFB9* are similar genes from the recently expanded cluster D on chromosome 27, and their genomic locations overlap (Table 1*). DEFB15* and *DEFB106A* also map to cluster D, and also overlap (Table 1). Using a lower threshold of r=0.85 identifies *DEFB113, DEFB112* to *DEFB110* as a possible copy number variable block on cluster C. However, full resolution of breakpoints of these complex and highly variable regions will require long-read sequencing and multiple complete genome assemblies.

We analysed β-defensin CNV in Holstein and Jersey breed cattle from the national Irish dairy herd by developing a ddPCR for a subset of β-defensin genes, selected for their likely importance in fertility. After validating assay accuracy using matched WGS data from the same individual (Supplementary figure 3), and validating assay precision using repeat testing, we typed 20 Holstein bulls and 15 Jersey bulls (Supplementary figure 4). We confirmed CNV of the same range in these cattle, including the extensive multiallelic CNV of *DEFB103*. This shows that β-defensin genes are extensively copy number variable in the Irish national dairy herd, and may be responsible for phenotypic variation across individuals.

### Expression of β-defensins in testis from calf and adult

Given previous evidence that most β-defensins are expressed in the testis, we wished to examine this in cattle in detail across our catalogue of β-defensins. We decided to use published RNASeq data from 3 (13-week-old) calves and 7 adult bulls (3 and 9 years and 11 months old) to perform a focused analysis of β-defensin expression in the testis during sexual maturation. By quantifying transcript levels, expression of 27 β-defensin genes was detected out of 44 (supplementary table 6), with 14 showing a higher expression (FDR<0.1) in adult testis compared to calf testis in Hereford bulls (Table 2).

**Table 2.**
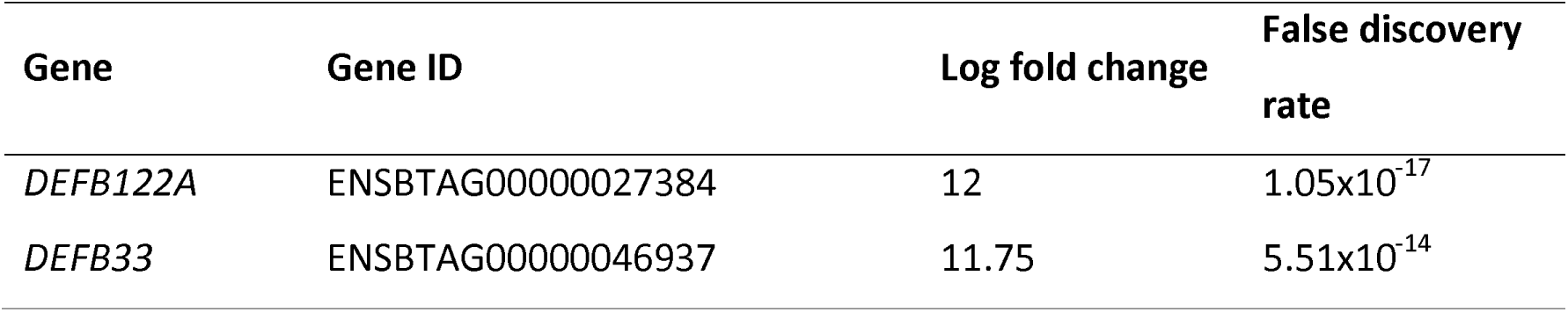
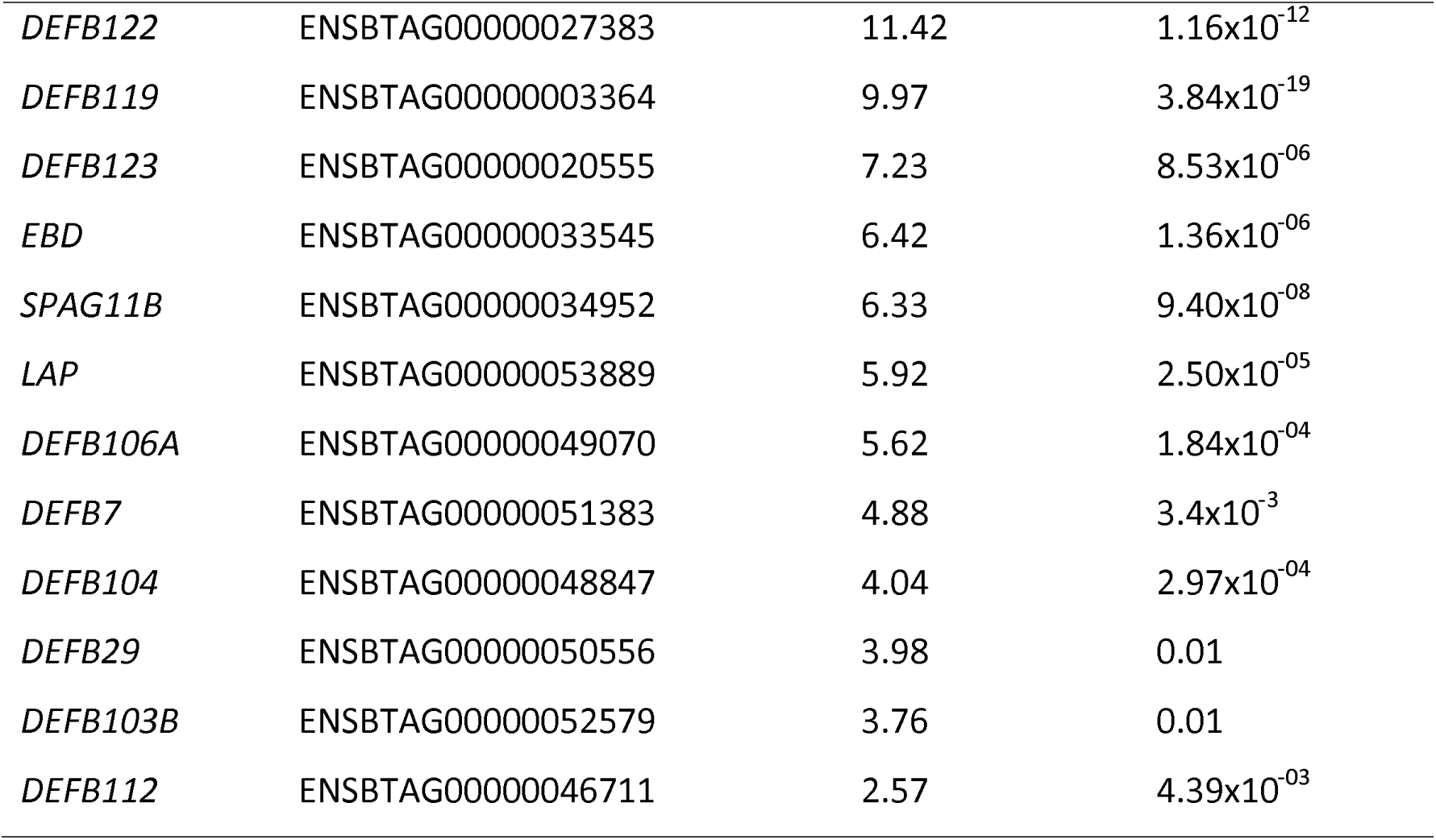
β-befensin genes upregulated in adult bull testes. Gene Gene ID Log fold change False discovery rate.

Previous work has shown that some β-defensins, including those involved in fertility, show an extended C-terminal tail with the potential to be heavily glycosylated, and to form part of the sperm glycocalyx, for example β-defensin 126 (Narciandi et al., 2016). We analysed the 14 β-defensin predicted protein sequences whose genes show an increase in expression at sexual maturity for presence of an extended C-terminal tail and glycosylation potential (Supplementary table 7). Of the 14, two (BBD119, SPAG11B) showed long C-terminal tails with at least potential O-glycosylation site, four (BBD122, BBD122A, BBD104, BBD29) showed a long-terminal tail with no glycosylation sites, and eight (BBD123, BBD112, BBD33, BBD15, BBD7, BBD103, EBD and LAP) showed no long C-terminal tail. The diversity of predicted structures of the upregulated β-defensins suggests that they have a variety of roles in the testis, potentially including glycocalyx formation and direct antimicrobial action.

### Expression of β-defensin in caput region of epididymis

The epididymis is a complex organ, and any expression data based on nucleic extraction from the whole organ will not be able to distinguish expression in different cell types which can limit interpretation of expression levels of particular genes. To address this in part, we used RNASeq data from the caput of the epididymis of 94 bulls to assess the transcript levels of 44 β-defensin genes (Figure 2). All 44 β-defensins are expressed to a greater or lesser extent in the caput, with, for example, *DEFB124, DEFB29* expressed at high levels and *TAP* and *DEFB118* expressed at low levels. Indeed, a previous analysis of this data found that four β-defensins (*DEFB110, DEFB124, DEFB121 and DEFB114*) were among the fifty most-tissue specific/tissue enriched transcripts in the epididymis (Mapel et al., 2024). However, all expressed defensins show a high degree of variation between different bulls for any particular β-defensin, often over three orders of magnitude. For some genes, such as *DEFB134* and *DEFB402*, this range includes several bulls who show no expression of the gene at all.

**Figure 2.**
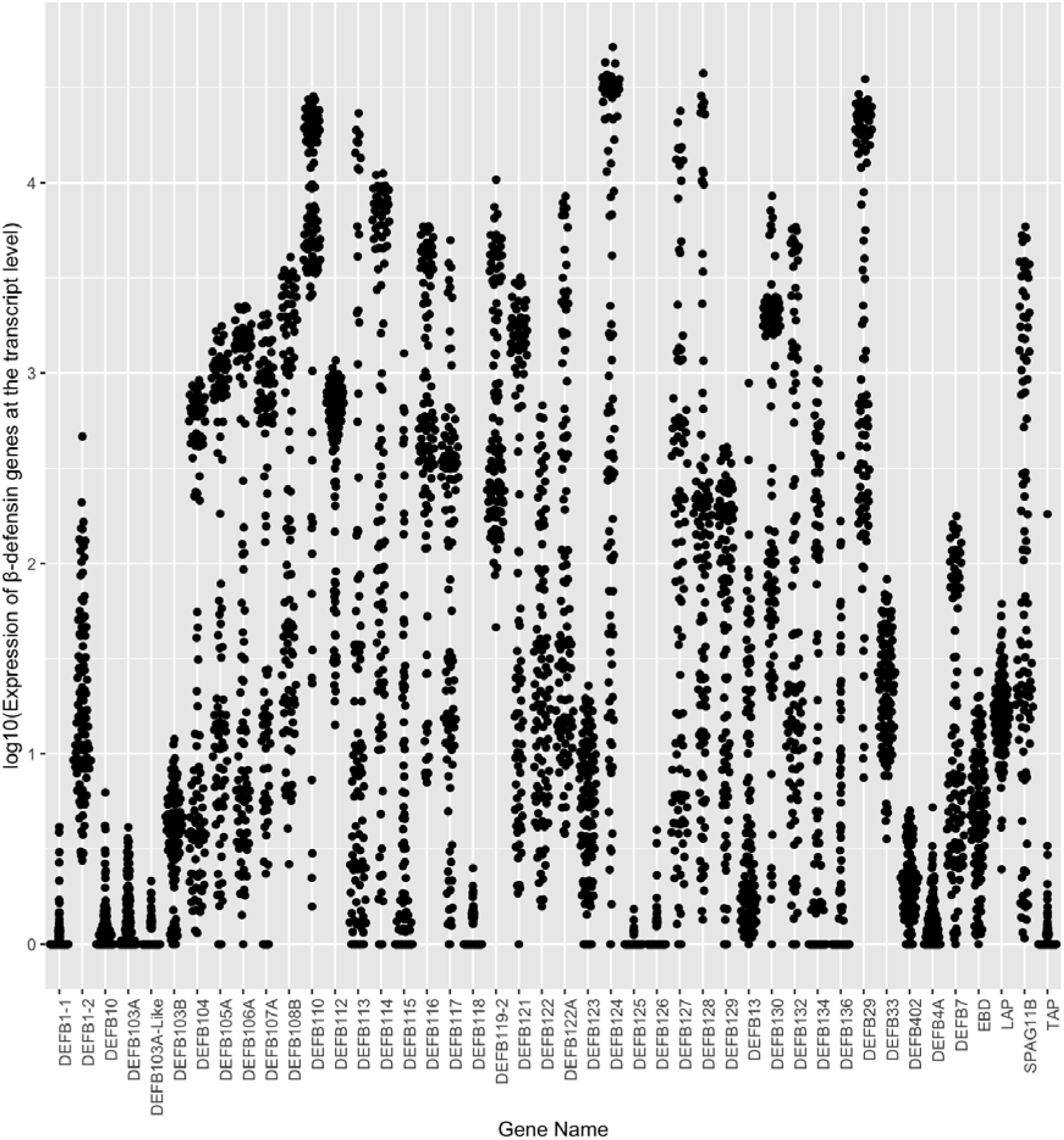
Expression of bovine β-defensin genes in the caput of the epidydimis.

It is likely that the pattern of β-defensin gene expression changes along the epididymis from caput to cauda. Strong regional expression of β-defensins in the epididymis has been shown in cattle (Narciandi et al., 2011) and rats (Dorin and Barratt, 2014). *DEFB103* is strongly expressed in the cauda of cattle epididymis (as is its orthologue *Defb14* in rats), as shown by RT-PCR (Patil et al., 2005), but our analysis shows that it is weakly expressed in the caput. Therefore, absence or low expression of a particular gene in the caput does not reflect on expression levels elsewhere in the epididymis, nor on its potential role in fertility.

Nevertheless, the extensive variation of expression of the defensins could potentially have consequences on fertility and reproduction-associated phenotypes.

### Correlation of gene expression with gene copy number

The individual variation in expression level will be due to a mixture of genetic, environmental and possibly technical factors. Given the extensive CNV of these genes, this is a strong candidate to explain at least part of any genetic variation contributing to the variation in expression levels between individuals. This could be through a gene dosage effect, where an increase in the number of copies of a gene leads to a concomitant increase in transcript level, or through more complex effects such as disrupting expression levels through higher-order chromatin structure at the locus.

Correlating the expression levels of the 44 genes against estimates of gene CNV identified four genes with a nominal p value <0.05 where expression was correlated with gene copy number (Supplementary table 7, Figure 3). For *DEFB1-1*, examination of the data shows that the correlation is driven by outlying higher expression levels in an overall very low-expressed gene. However, for the remaining three genes, *DEFB10* shows positively correlation but two (*DEFB127* and *DEFB103*) show negative correlation. The positive correlation can be explained at least in part by a gene dosage effect, and given that the protein encoded by *DEFB10*, BBD10, is part of the recently duplicated clade including LAP and TAP, is upregulated by vitamin D and is likely to be antimicrobial (Kweh et al., 2019), although this has not been shown directly. *DEFB103* orthologues have been shown to be antimicrobial but also involved in cell signalling (Harvey et al., 2013). *DEFB127* has a long C-terminal tail (45 amino acids after last cysteine), 4 predicted glycosylation sites, and is likely to be involved in fertility by binding to sperm directly.

**Figure 3.**
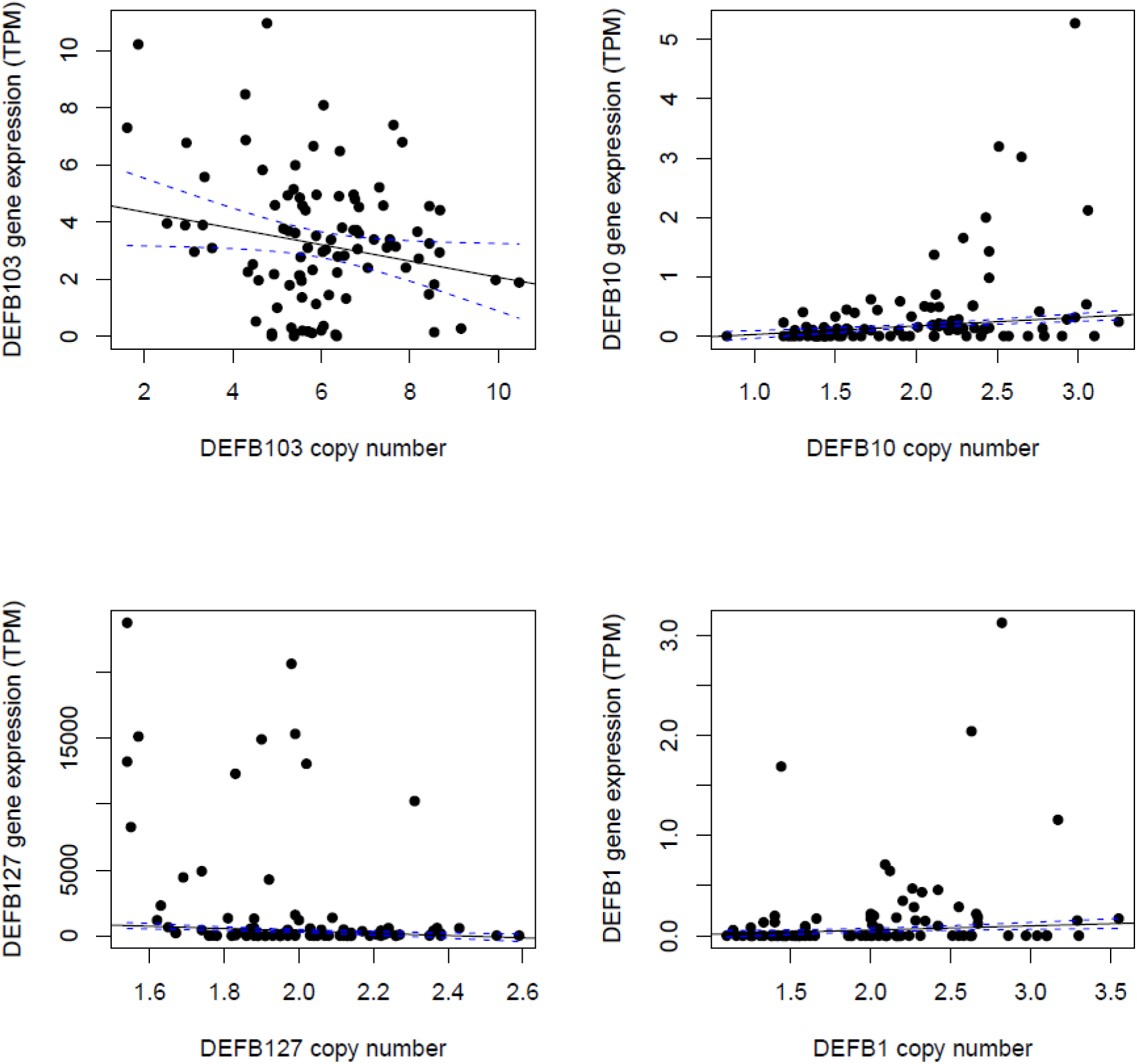
Copy number variation and gene expression of selected β-defensins. DEFB10 (a), DEFB127 (b), DEFB1-1 (c), and DEFB103 (d) genes shown. The X-axis represents the diploid copy number, while the Y-axis indicates the expression of the gene in transcripts per million.

## Discussion

In this study, we have fully characterised the complement of β-defensin genes in cattle, and have constructed a phylogenetic tree to identify their human orthologues, and β-defensin family expansions in the bovine lineage. This builds on previous work, starting with identification of the genes, including *LAP* and *TAP*, that we know are a recent gene family expansion on chromosome 13 (Roosen et al., 2004). These were shown initially to be expressed in udders. A more complete search of the cattle genome assembly UMD 3.1 identified 57 β-defensin open-reading-frames, of which 53 were full-length (Cormican et al., 2008). In this study, by using the recent high quality ARS-UCD1.2 genome assembly, we refine this complement to 55 full-length beta defensin genes, and identify 1:1 human orthologues for 35 bovine β-defensin genes, suggesting conservation of function of these β-defensins for ∼90 million years. In contrast, the cluster of β-defensins on chromosome 13 have evolved by more recent duplication and divergence. This chromosome 13 β-defensin cluster generally have a broader expression profile across tissues compared to older β- defensins, which are generally restricted, or most highly expressed, in the testis (Cormican et al., 2008). The cluster includes *LAP* and *TAP*, encoding for lingual antimicrobial peptide and tracheal antimicrobial peptide respectively, and these two proteins have been shown to be directly antimicrobial, and involved in the immune response to various disease.

Orthologues of at least some of these genes have been identified in sheep (Hall et al., 2017; Huttner et al., 1998), goat and water buffalo (as reviewed by Meade et al. (2014))

We also show that genes encoding three different predicted BBD103 proteins map to three different regions on the ARS-UCD1.2 genome assembly. BBD103B is short, and most similar to human β-defensin 103, suggesting that it is its orthologue. BBD103A has a 14-amino acid segment at the N-terminus, which is likely to disrupt any signal sequencing processing and consequent secretion, and a long C-terminal tail. Compared to BBD103B BBD103-like does not have a predicted initial methionine. It is likely that only BBD103B is a functional protein, however, all three transcripts are detected in the caput of the epididymis, and the CNV we observe for *DEFB103* does not distinguish between the three related genes. Multiple copies of the *DEFB103* gene are likely to encode a mixture of the three related transcripts: A, A-like and B, but analysing the diversity within the CNV requires further investigation. Our observation of *DEFB103* CNV might explain in part the observations in previous publications on this gene – Mirabzadeh-Ardakani et al (2014) analyse *DEFB103B*, but document extra copies of *DEFB103* in earlier genome assemblies (Mirabzadeh-Ardakani et al., 2014).

The genes on clustered bovine chromosome 8 - *DEFB136*, *DEFB134*, and *DEFB43* (also known as *DEFB131*) do not form an evolutionary cluster, but share apparent evolutionary relationships with genes on other chromosomes. They preserve synteny, but in inverted order, with their orthologues in the sheep genome (Hall et al., 2017), synteny with their orthologues in dogs (Patil et al., 2005), and are in a highly structurally variable region in humans. A full analysis of β-defensin evolution in artiodactyls will clarify the evolution of this cluster, and the chromosome 13 cluster.

Previous work has suggested that many beta defensins show CNV across cattle breeds (Bickhart et al., 2012). However, although existence of CNV was extensively validated, estimates of copy number were high, on five cattle with low sequencing coverage, shorter read lengths and were based on an early cattle genome assembly. We took a gene-focused approach to investigate CNV in 191 cattle across seven breeds from the 1000 bulls project and previously published data, and confirmed/extended our analysis into the Irish national herd of Holstein and Jersey dairy bulls. We found extensive CNV of β-defensin genes across all breeds, particularly the Holstein-Friesians. We show that certain genes covary, suggesting that in some cases there are multiple genes on a single copy number variable genomic segment. However, because of the extensive variability between breeds and individuals, refining the breakpoints of these CNV regions will remain a challenge until multiple telomere-to-telomere reference genome assemblies are available across breeds.

β-defensins are expressed in the epididymis, and are likely to be important for fertility and reproduction by maintaining healthy, functioning sperm, either directly by being a component of the glycocalyx, for example, or indirectly by providing an innate immune defence against microbes present in the female reproductive tract (Fernandez-Fuertes et al., 2016; Ganz, 2004; Yudin et al., 2008). Defensins that show increased expression in adult testis compared to calf testis are likely to have been upregulated during sexual maturation and therefore be particularly important in the function of the sperm. The defensins found to be upregulated include both highly conserved testis specific defensins with long, potentially glycosylated, C-terminal tails, and more recent defensins, shorter, with known antimicrobial and immunomodulatory functions. Of the 14 found to be upregulated in this study, in a study on selected β-defensins, *DEFB119* had been found by RT-PCR previously to be absent in calf testis but highly expressed in adult testis (Narciandi et al., 2011). Taken together, our data suggests defensins with a variety of functional roles are important in fertility, and highlights a selected number to be prioritised for further functional analysis.

Sperm maturation occurs in the epididymis is divided into cauda, caput and corpus. It has been previously shown that defensins are expressed in the epididymis of bulls, with region-specific expression shown, at least in rodents. In this study, we show that almost all β-defensins are expressed in the caput, but show extensive inter-individual variation. A variety of factors may influence expression levels, including age, time and season of sampling, and emphasises that multiple individuals need to be included in an experiment in order to make secure inferences about tissue- and region-specific expression.

Genetic variation can influence expression levels of a gene, and CNV of β-defensins has been associated with expression levels in humans (James et al., 2018, 2020; Janssens et al., 2010). We used the caput expression data to test whether gene copy number accounted for at least part of the expression variation observed between individual bulls. We found that, for four genes, expression level was correlated with copy number, although accounting for a small fraction (<14%) of the variation. Interestingly, for two of the genes the association was negative, suggesting that these effects might be through alterations regulation/distance from an enhancer, or alterations in chromatin structure. The absence of an effect on copy number on expression levels of other genes suggests that extra copies of these genes may be silenced, at least in the epididymis, or the effect on expression is far outweighed by other genetic or environmental factors.

Taken together, we elucidate the evolutionary history and demonstrate extensive CNV of cattle β-defensins, and highlight several genes that are strong candidates for a functional effect of CNV on fertility. In particular, *DEFB103* is expressed in the epididymis caput, shows extensive multiallelic CNV which correlates with its expression level, and is upregulated on sexual maturity. Functional characterisation of the role of *DEFB103* and the encoded protein BBD103 in reproduction and fertility is a priority.

## Funding

OS was funded by the Turkish Ministry of National Education, Republic of Turkiye postgraduate study abroad program and a BBSRC Impact Accelerator Award.

JO was funded by PhD studentship as part of the Wellcome Trust Genetic Epidemiology and Public Health Genomics Doctoral Training Programme by grant number 218505/Z/19/Z.

SF was funded by the Marie Skłodowska-Curie Doctoral Network, BullNet. Grant number 101120104.

## Acknowledgements

This research used the ALICE High Performance Computing Facility at the University of Leicester. We are grateful to Dovea Genetics, Thurlus, Ireland for providing availability of positive control bull semen straws. We thank Maymun Jama for advice on ddPCR, and Diana Martin for technical support.

## Authorship statement

Ozge Sidekli: Conceptualisation, methodology, investigation, writing – review and editing, visualisation, formal analysis, data curation, funding acquisition

John Oketch: Formal analysis, data curation

Sean Fair: Resources

Kieran Meade: Conceptualisation, funding acquisition, supervision, resources, writing – review and editing.

Edward Hollox: Conceptualisation, funding acquisition, supervision, writing - original draft, visualisation, project administration.

## Data availability statement

European Nucleotide Archive (https://www.ebi.ac.uk/ena) Accession numbers for DNA sequencing data are given in supplementary tables 1, 2, 3 and 6.

Raw gene copy number estimates by ddPCR and WGS are freely available at Leicester Research Archive https://doi.org/10.25392/leicester.data.25992484.v1 and https://doi.org/10.25392/leicester.data.25991863.v1.

---

**Supplementary Table 1.** Cattle from the 1000 bulls consortium for whole genome sequence (WGS) analysis.

| Study Accession | Bull ID | Run Accession | Breed | Coverage (x) |
| --- | --- | --- | --- | --- |
| PRJNA474946 | HERIRLM_IMAG_000047 | SRR7363649 | Hereford | 11.3 |
| PRJNA474946 | HERIRLM_IMAG_000015 | SRR7363653 | Hereford | 10.4 |
| PRJNA474946 | HERGBRM000HPBP38128 | SRR7363655 | Hereford | 13.4 |
| PRJNA474946 | HERIRLM131456220077 | SRR7363658 | Hereford | 17.7 |
| PRJNA474946 | HERIRLM_IMAG_000052 | SRR7363667 | Hereford | 13.2 |
| PRJNA474946 | HERIRLM371254460275 | SRR7363678 | Hereford | 15.6 |
| PRJNA474946 | CHAIRLM00000KHTL001 | SRR7363652 | Charolais | 17.9 |
| PRJNA474946 | CHAFRAM007121319604 | SRR7363748 | Charolais | 23.6 |
| PRJNA474946 | CHAFRAM007984101105 | SRR7363750 | Charolais | 14 |
| PRJNA474946 | CHAFRAM008590100065 | SRR7363761 | Charolais | 18.2 |
| PRJNA474946 | CHAFRAM008526937032 | SRR7363763 | Charolais | 19.5 |
| PRJNA474946 | CHAFRAM008532799816 | SRR7363764 | Charolais | 15.6 |
| PRJNA474946 | CHAFRAM008526936895 | SRR7363765 | Charolais | 11.2 |
| PRJNA474946 | CHAFRAM008526936641 | SRR7363766 | Charolais | 14.8 |
| PRJNA474946 | CHAFRAM008526867944 | SRR7363767 | Charolais | 11.4 |
| PRJNA474946 | CHAFRAM008592106619 | SRR7363771 | Charolais | 17.1 |
| PRJNA474946 | CHAIRLM000000MJH014 | SRR7363780 | Charolais | 11.7 |
| PRJNA474946 | CHAIRLM000000WXD001 | SRR7363781 | Charolais | 11.6 |
| PRJNA474946 | CHAIRLM000000RRB002 | SRR7363782 | Charolais | 16.5 |
| PRJNA474946 | CHAFRAM008690101065 | SRR7363773 | Charolais | 12.2 |
| PRJNA474946 | CHAGBRM000MF0034726 | SRR7363774 | Charolais | 21.2 |
| PRJNA474946 | CHAGBRM000FF0013463 | SRR7363775 | Charolais | 10.7 |
| PRJNA474946 | CHAIRLM000000AGU001 | SRR7363776 | Charolais | 11.7 |
| PRJNA474946 | CHAGBRM009587367887 | SRR7363777 | Charolais | 13.7 |
| PRJNA474946 | CHAIRLM000000BIC001 | SRR7363778 | Charolais | 12.5 |
| PRJNA474946 | CHAIRLM000000HDHM001 | SRR7363654 | Charolais | 17.3 |
| PRJNA474946 | CHAIRLM000000HNRN006 | SRR7363656 | Charolais | 13.3 |
| PRJNA474946 | CHAIRLM000000GHFI004 | SRR7363657 | Charolais | 21.1 |
| PRJNA474946 | CHAFRAM000310249639 | SRR7363727 | Charolais | 10 |
| PRJNA474946 | CHAFRAM005815216302 | SRR7363731 | Charolais | 13 |
| PRJNA474946 | CHAFRAM008596102216 | SRR7363769 | Charolais | 16.2 |
| PRJNA474946 | CHAFRAM005898106607 | SRR7363743 | Charolais | 13.5 |
| PRJNA474946 | LIMFRAM001692111209 | SRR7363772 | Limousin | 11.3 |
| PRJNA474946 | LIMFRAM001931738155 | SRR7363783 | Limousin | 17.8 |
| PRJNA474946 | LIMFRAM001997013395 | SRR7363784 | Limousin | 16.4 |
| PRJNA474946 | LIMFRAM001931107913 | SRR7363786 | Limousin | 18.5 |
| PRJNA474946 | LIMGBRM000000GEO2020 | SRR7363702 | Limousin | 11.1 |
| PRJNA474946 | LIMFRAM001930818375 | SRR7363785 | Limousin | 11 |
| PRJNA474946 | LIMFRAM003683001518 | SRR7363711 | Limousin | 17.1 |
| PRJNA474946 | LIMFRAM005500194039 | SRR7363740 | Limousin | 12.5 |
| PRJNA474946 | LIMIRLM000000DERA004 | SRR7363713 | Limousin | 12.9 |
| PRJNA474946 | LIMIRLM000000GVO3176 | SRR7363719 | Limousin | 11.2 |
| PRJNA474946 | LIMIRLM000000LNCJ002 | SRR7363717 | Limousin | 17.8 |
| PRJNA474946 | LIMIRLM000000PELA004 | SRR7363716 | Limousin | 11 |
| PRJNA474946 | LIMIRLM281084560407 | SRR7363720 | Limousin | 14.2 |
| PRJNA474946 | LIMIRLM281523440569 | SRR7363682 | Limousin | 12.2 |
| PRJNA474946 | LIMIRLM000000PELC017 | SRR7363721 | Limousin | 10.1 |
| PRJNA474946 | LIMFRAM008789003682 | SRR7363708 | Limousin | 10.9 |
| PRJNA474946 | LIMGBRM000000MDS03012 | SRR7363714 | Limousin | 11.7 |
| PRJNA474946 | LIMIRLM00000KGEH003 | SRR7363718 | Limousin | 14.7 |
| PRJNA474946 | LIMFRAM003693000206 | SRR7363704 | Limousin | 10.8 |
| PRJNA474946 | SIMIRLM_IIMAG_000075 | SRR7363755 | Simmental | 12.2 |
| PRJNA474946 | SIMIRLM_IIMAG_000046 | SRR7363756 | Simmental | 18.6 |
| PRJNA474946 | SIMGBRM00000M001746 | SRR7363757 | Simmental | 14.1 |
| PRJNA474946 | SIMGBRM00000I000563 | SRR7363758 | Simmental | 18.4 |
| PRJNA474946 | SIMAUTM0000871580242 | SRR7363683 | Simmental | 10.8 |
| PRJNA474946 | SIMGBRM00000I000139 | SRR7363684 | Simmental | 11.5 |
| PRJNA474946 | SIMGBRM00000I000377 | SRR7363690 | Simmental | 10.3 |
| PRJNA474946 | SIMAUTM000000620572 | SRR7363685 | Simmental | 15.8 |
| PRJNA474946 | SIMAUTM000212220633 | SRR7363687 | Simmental | 13.2 |
| PRJNA474946 | AANGBRM00000IUN225 | SRR7363670 | Angus | 14.3 |
| PRJNA474946 | AANGBRM000000785793 | SRR7363671 | Angus | 12.5 |
| PRJNA474946 | AANGBRM00000ELFP126 | SRR7363672 | Angus | 12.4 |
| PRJNA474946 | AANGBRM00000IFPH318 | SRR7363673 | Angus | 12.8 |
| PRJNA474946 | AANGBRM00000GCAR117 | SRR7363669 | Angus | 13.3 |
| PRJNA474946 | AANGBRM000000880440 | SRR7363674 | Angus | 10.4 |
| PRJNA474946 | AANGBRM00000MEPV221 | SRR7363675 | Angus | 11.8 |
| PRJNA474946 | AANGBRM000020020052 | SRR7363677 | Angus | 12.9 |
| PRJNA474946 | AANGBRM0000MQUO1115 | SRR7363696 | Angus | 12.9 |
| PRJNA474946 | AANGBRM000020041290 | SRR7363706 | Angus | 11.3 |
| PRJEB9343 | INRA_HOL0002 | ERR883661 | Holstein | 12.6 |
| PRJEB9343 | INRA_HOL0003 | ERR883662 | Holstein | 18.8 |
| PRJEB9343 | INRA_HOL0004 | ERR883663 | Holstein | 17.3 |
| PRJEB9343 | INRA_HOL0005 | ERR883664 | Holstein | 18.8 |
| PRJEB9343 | INRA_HOL0006 | ERR883665 | Holstein | 11.4 |
| PRJEB9343 | INRA_HOL0007 | ERR883666 | Holstein | 18.8 |
| PRJEB9343 | INRA_HOL0008 | ERR883667 | Holstein | 14.1 |
| PRJEB9343 | INRA_HOL0009 | ERR883668 | Holstein | 11.1 |
| PRJEB9343 | INRA_HOL0010 | ERR883669 | Holstein | 19.3 |
| PRJEB9343 | INRA_HOL0011 | ERR883670 | Holstein | 11.2 |
| PRJEB9343 | INRA_HOL0013 | ERR883672 | Holstein | 15.6 |
| PRJEB9343 | INRA_HOL0015 | ERR883674 | Holstein | 12.9 |
| PRJEB9343 | INRA_HOL0016 | ERR883675 | Holstein | 11.8 |
| PRJEB9343 | INRA_HOL0017 | ERR883676 | Holstein | 11.1 |
| PRJEB9343 | INRA_HOL0018 | ERR883677 | Holstein | 11.3 |
| PRJEB9343 | INRA_HOL0021 | ERR883680 | Holstein | 12.5 |
| PRJEB9343 | INRA_HOL0022 | ERR883681 | Holstein | 11.2 |
| PRJEB9343 | INRA_HOL0023 | ERR883682 | Holstein | 11.3 |
| PRJEB9343 | INRA_HOL0024 | ERR883683 | Holstein | 14.4 |
| PRJEB9343 | INRA_HOL0026 | ERR883685 | Holstein | 15.8 |
| PRJEB9343 | INRA_HOL0027 | ERR883686 | Holstein | 14.2 |
| PRJEB18113 | RM666 | ERR1746312 | Holstein | 11.5 |
| PRJEB18113 | RM767 | ERR1746323 | Holstein | 15 |
| PRJEB18113 | RM774 | ERR1746325 | Holstein | 14.7 |
| PRJNA238491 | Btau_HOL_0007 | SRR1262661 | Holstein | 8.2 |
| PRJNA238491 | Btau_HOL_0009 | SRR1262663 | Holstein | 7.6 |
| PRJNA238491 | Btau_HOL_0014 | SRR1262668 | Holstein | 9.8 |
| PRJNA238491 | Btau_HOL_0018 | SRR1262672 | Holstein | 11.8 |
| PRJNA238491 | Btau_HOL_0037 | SRR1262691 | Holstein | 10.2 |
| PRJNA238491 | Btau_HOL_0048 | SRR1262702 | Holstein | 18.1 |

**Supplementary table 2.** Cattle used for RNAseq analysis.

| Accession number | Sample Name | Breed | Biosample | Age | Age group |
| --- | --- | --- | --- | --- | --- |
| PRJNA760322 | SRR15711831 | Hereford | Testis | ~13 weeks | Calf |
| PRJNA760322 | SRR15711832 | Hereford | Testis | ~13 weeks | Calf |
| PRJNA760322 | SRR15711833 | Hereford | Testis | ~13 weeks | Calf |
| PRJNA760322 | SRR15711834 | Hereford | Testis | 3 years | Adult |
| PRJNA760322 | SRR15711835 | Hereford | Testis | 3 years | Adult |
| PRJNA760322 | SRR15711836 | Hereford | Testis | 3 years | Adult |
| PRJNA263600 | SRR1640128 | Hereford | Testis | 9 years | Adult |
| PRJNA379574 | SRR5363137 | Hereford | Testis | 9 years | Adult |
| PRJNA526257 | SRR8703147 | Hereford | Testis | 11 month | Adult |
| PRJNA526257 | SRR8703148 | Hereford | Testis | 11 month | Adult |

**Supplementary table 3.** The details of the DNA sequencing (paired-end; 150bp) samples obtained from testis and RNA sequencing (paired-end; 150bp) samples obtained from caput epididymis.

| DNA Sequence (paired-end; 150bp)<br>samples from Testis |  |  |  | RNA sequence (paired-end; 150bp) |  |
| --- | --- | --- | --- | --- | --- |
| Study Accession | Sample Accession | Run Accession | Coverage | Study Accession | Sample Acc |
| PRJEB46995 | SAMEA111328912 | ERR10164310 | 12.51 | PRJEB46995 | SAMEA1136 |
| PRJEB46995 | SAMEA111328913 | ERR10164311 | 15.34 | PRJEB46995 | SAMEA1136 |
| PRJEB46995 | SAMEA111328914 | ERR10164312 | 15.14 | PRJEB46995 | SAMEA1136 |
| PRJEB46995 | SAMEA111328916 | ERR10164314 | 13.12 | PRJEB46995 | SAMEA1136 |
| PRJEB46995 | SAMEA111328917 | ERR10164315 | 16.81 | PRJEB46995 | SAMEA1136 |
| PRJEB46995 | SAMEA111328919 | ERR10164317 | 14.51 | PRJEB46995 | SAMEA1136 |
| PRJEB46995 | SAMEA111328920 | ERR10164318 | 15.42 | PRJEB46995 | SAMEA1136 |
| PRJEB46995 | SAMEA111328921 | ERR10164319 | 10.66 | PRJEB46995 | SAMEA1136 |
| PRJEB46995 | SAMEA111328922 | ERR10164320 | 20.76 | PRJEB46995 | SAMEA1136 |
| PRJEB46995 | SAMEA111328923 | ERR10164321 | 19.24 | PRJEB46995 | SAMEA1136 |
| PRJEB46995 | SAMEA111328924 | ERR10164322 | 13.95 | PRJEB46995 | SAMEA1136 |
| PRJEB46995 | SAMEA111328925 | ERR10164323 | 16.66 | PRJEB46995 | SAMEA1136 |
| PRJEB46995 | SAMEA111328926 | ERR10164324 | 13.78 | PRJEB46995 | SAMEA1136 |
| PRJEB46995 | SAMEA111328928 | ERR10164326 | 11.97 | PRJEB46995 | SAMEA1136 |
| PRJEB46995 | SAMEA111328929 | ERR10164327 | 11.37 | PRJEB46995 | SAMEA1136 |
| PRJEB46995 | SAMEA111328930 | ERR10164328 | - | PRJEB46995 | SAMEA1136 |
| PRJEB46995 | SAMEA111328931 | ERR10164329 | 14.24 | PRJEB46995 | SAMEA1136 |
| PRJEB46995 | SAMEA111328932 | ERR10164330 | 15 | PRJEB46995 | SAMEA1136 |
| PRJEB46995 | SAMEA111328933 | ERR10164331 | 10.71 | PRJEB46995 | SAMEA1136 |
| PRJEB46995 | SAMEA111328934 | ERR10164332 | 10.23 | PRJEB46995 | SAMEA1136 |
| PRJEB46995 | SAMEA111328935 | ERR10164333 | 10.13 | PRJEB46995 | SAMEA1136 |
| PRJEB46995 | SAMEA111328938 | ERR10164336 | 11.69 | PRJEB46995 | SAMEA1136 |
| PRJEB46995 | SAMEA111328940 | ERR10164338 | - | PRJEB46995 | SAMEA1136 |
| PRJEB46995 | SAMEA111328941 | ERR10164339 | 14.63 | PRJEB46995 | SAMEA1136 |
| PRJEB46995 | SAMEA111328942 | ERR10164340 | 11.18 | PRJEB46995 | SAMEA1136 |
| PRJEB46995 | SAMEA111328943 | ERR10164341 | 12.44 | PRJEB46995 | SAMEA1136 |
| PRJEB28191 | SAMEA9539940 | ERR6501753 | 15.09 | PRJEB46995 | SAMEA1136 |
| PRJEB28191 | SAMEA9539941 | ERR6501754 | 10.66 | PRJEB46995 | SAMEA1136 |
| PRJEB28191 | SAMEA9539943 | ERR6501756 | 27.04 | PRJEB46995 | SAMEA1136 |
| PRJEB28191 | SAMEA9539945 | ERR6501758 | 16.4 | PRJEB46995 | SAMEA1136 |
| PRJEB28191 | SAMEA9539946 | ERR6501759 | 12.55 | PRJEB46995 | SAMEA1136 |
| PRJEB28191 | SAMEA9539947 | ERR6501760 | 12.85 | PRJEB46995 | SAMEA1136 |
| PRJEB28191 | SAMEA9539948 | ERR6501761 | 13.99 | PRJEB46995 | SAMEA1136 |
| PRJEB28191 | SAMEA9539949 | ERR6501762 | 11 | PRJEB46995 | SAMEA1136 |
| PRJEB28191 | SAMEA9539950 | ERR6501763 | 32.31 | PRJEB46995 | SAMEA1136 |
| PRJEB28191 | SAMEA9539952 | ERR6501765 | 27.37 | PRJEB46995 | SAMEA1136 |
| PRJEB28191 | SAMEA9539953 | ERR6501766 | 15.68 | PRJEB46995 | SAMEA1136 |
| PRJEB28191 | SAMEA9539954 | ERR6501767 | 11.52 | PRJEB46995 | SAMEA1136 |
| PRJEB28191 | SAMEA9539955 | ERR6501768 | 10.42 | PRJEB46995 | SAMEA1136 |
| PRJEB28191 | SAMEA9539956 | ERR6501769 | 12.12 | PRJEB46995 | SAMEA1136 |
| PRJEB28191 | SAMEA9539957 | ERR6501770 | 20.32 | PRJEB46995 | SAMEA1136 |
| PRJEB28191 | SAMEA9539958 | ERR6501771 | 21.06 | PRJEB46995 | SAMEA1136 |
| PRJEB28191 | SAMEA9539959 | ERR6501772 | 15.12 | PRJEB46995 | SAMEA1136 |
| PRJEB28191 | SAMEA9539961 | ERR6501774 | 13.25 | PRJEB46995 | SAMEA1136 |
| PRJEB28191 | SAMEA9539962 | ERR6501775 | 13.87 | PRJEB46995 | SAMEA1136 |
| PRJEB28191 | SAMEA9539963 | ERR6501776 | 17.69 | PRJEB46995 | SAMEA1136 |
| PRJEB28191 | SAMEA9539964 | ERR6501777 | 16.27 | PRJEB46995 | SAMEA1136 |
| PRJEB28191 | SAMEA9539965 | ERR6501778 | 11.13 | PRJEB46995 | SAMEA1136 |
| PRJEB28191 | SAMEA9539966 | ERR6501779 | 16.46 | PRJEB46995 | SAMEA1136 |
| PRJEB28191 | SAMEA9539967 | ERR6501780 | 12.27 | PRJEB46995 | SAMEA1136 |
| PRJEB28191 | SAMEA9539968 | ERR6501781 | 23.62 | PRJEB46995 | SAMEA1136 |
| PRJEB28191 | SAMEA9539969 | ERR6501782 | 13.49 | PRJEB46995 | SAMEA1136 |
| PRJEB28191 | SAMEA9539970 | ERR6501783 | 20.56 | PRJEB46995 | SAMEA1136 |
| PRJEB28191 | SAMEA9539971 | ERR6501784 | 14.86 | PRJEB46995 | SAMEA1136 |
| PRJEB28191 | SAMEA9539972 | ERR6501785 | 20.02 | PRJEB46995 | SAMEA1136 |
| PRJEB28191 | SAMEA9539973 | ERR6501786 | 13.91 | PRJEB46995 | SAMEA1136 |
| PRJEB28191 | SAMEA9539974 | ERR6501787 | 9.87 | PRJEB46995 | SAMEA1136 |
| PRJEB28191 | SAMEA9539975 | ERR6501788 | 12.72 | PRJEB46995 | SAMEA1136 |
| PRJEB28191 | SAMEA9539976 | ERR6501789 | 15.04 | PRJEB46995 | SAMEA1136 |
| PRJEB28191 | SAMEA9539977 | ERR6501790 | 14.4 | PRJEB46995 | SAMEA1136 |
| PRJEB28191 | SAMEA9539978 | ERR6501791 | 13.85 | PRJEB46995 | SAMEA1136 |
| PRJEB28191 | SAMEA9539979 | ERR6501792 | 13.02 | PRJEB46995 | SAMEA1136 |
| PRJEB28191 | SAMEA9539980 | ERR6501793 | 12.22 | PRJEB46995 | SAMEA1136 |
| PRJEB28191 | SAMEA9539981 | ERR6501794 | 9.11 | PRJEB46995 | SAMEA1136 |
| PRJEB28191 | SAMEA9539982 | ERR6501795 | 10.62 | PRJEB46995 | SAMEA1136 |
| PRJEB28191 | SAMEA9539983 | ERR6501796 | 11.97 | PRJEB46995 | SAMEA1136 |
| PRJEB28191 | SAMEA9539984 | ERR6501797 | 15.56 | PRJEB46995 | SAMEA1136 |
| PRJEB28191 | SAMEA9539986 | ERR6501799 | 10.83 | PRJEB46995 | SAMEA1136 |
| PRJEB28191 | SAMEA9539987 | ERR6501800 | 16.39 | PRJEB46995 | SAMEA1136 |
| PRJEB28191 | SAMEA9539988 | ERR6501801 | 18.23 | PRJEB46995 | SAMEA1136 |
| PRJEB28191 | SAMEA9539989 | ERR6501802 | 17.39 | PRJEB46995 | SAMEA1136 |
| PRJEB28191 | SAMEA9539990 | ERR6501803 | 8.99 | PRJEB46995 | SAMEA1136 |
| PRJEB28191 | SAMEA9539991 | ERR6501804 | 13.19 | PRJEB46995 | SAMEA1136 |
| PRJEB28191 | SAMEA9539992 | ERR6501805 | 10.56 | PRJEB46995 | SAMEA1136 |
| PRJEB28191 | SAMEA9539993 | ERR6501806 | 7.85 | PRJEB46995 | SAMEA1136 |
| PRJEB28191 | SAMEA9539995 | ERR6501808 | 19.65 | PRJEB46995 | SAMEA1136 |
| PRJEB28191 | SAMEA9539996 | ERR6501809 | 12.15 | PRJEB46995 | SAMEA1136 |
| PRJEB28191 | SAMEA9539998 | ERR6501811 | 14.67 | PRJEB46995 | SAMEA1136 |
| PRJEB28191 | SAMEA9539999 | ERR6501812 | 16.25 | PRJEB46995 | SAMEA1136 |
| PRJEB28191 | SAMEA9540000 | ERR6501813 | 10.39 | PRJEB46995 | SAMEA1136 |
| PRJEB28191 | SAMEA9540001 | ERR6501814 | 8.74 | PRJEB46995 | SAMEA1136 |
| PRJEB28191 | SAMEA9540002 | ERR6501815 | 16.62 | PRJEB46995 | SAMEA1136 |
| PRJEB28191 | SAMEA9540003 | ERR6501816 | 31.77 | PRJEB46995 | SAMEA1136 |
| PRJEB28191 | SAMEA9540004 | ERR6501817 | 29.66 | PRJEB46995 | SAMEA1136 |
| PRJEB28191 | SAMEA9540005 | ERR6501818 | 15.07 | PRJEB46995 | SAMEA1136 |
| PRJEB28191 | SAMEA9540006 | ERR6501819 | 10.62 | PRJEB46995 | SAMEA1136 |
| PRJEB28191 | SAMEA9540007 | ERR6501820 | 9.78 | PRJEB46995 | SAMEA1136 |
| PRJEB28191 | SAMEA9540008 | ERR6501821 | 15.69 | PRJEB46995 | SAMEA1136 |
| PRJEB28191 | SAMEA9540009 | ERR6501822 | 12.74 | PRJEB46995 | SAMEA1136 |
| PRJEB28191 | SAMEA9540011 | ERR6501824 | 14.68 | PRJEB46995 | SAMEA1136 |
| PRJEB28191 | SAMEA9540013 | ERR6501826 | 19.27 | PRJEB46995 | SAMEA1136 |
| PRJEB28191 | SAMEA9540015 | ERR6501828 | 16.5 | PRJEB46995 | SAMEA1136 |
| PRJEB46995 | SAMEA113578975 | ERR11527327 | 15.09 | PRJEB46995 | SAMEA1136 |
| PRJEB46995 | SAMEA113578977 | ERR11527329 | 16.06 | PRJEB46995 | SAMEA1136 |
| PRJEB46995 | SAMEA113578978 | ERR11527330 | 13,42 | PRJEB46995 | SAMEA1136 |
| PRJEB46995 | SAMEA113578979 | ERR11527331 | 11.99 | PRJEB46995 | SAMEA1136 |
| PRJEB46995 | SAMEA113578980 | ERR11527332 | 12.7 | PRJEB46995 | SAMEA1136 |
| PRJEB46995 | SAMEA113578981 | ERR11527333 | 13.26 | PRJEB46995 | SAMEA1136 |
| PRJEB46995 | SAMEA113578967 | ERR11527319 | 8.98 | PRJEB46995 | SAMEA1136 |
| PRJEB46995 | SAMEA113578968 | ERR11527320 | 12.51 | PRJEB46995 | SAMEA1136 |
| PRJEB46995 | SAMEA113578969 | ERR11527321 | 15.14 | PRJEB46995 | SAMEA1136 |
| PRJEB46995 | SAMEA113578970 | ERR11527322 | 10.82 | PRJEB46995 | SAMEA1136 |
| PRJEB46995 | SAMEA113578971 | ERR11527323 | 12.2 | PRJEB46995 | SAMEA1136 |

**Supplementary Table 4.** Genomic location of 55 bovine β-defensin genes. Genomic locations are from the ARS-UCD1.genome assembly.

| Chromosome | Gene | Start | End |
| --- | --- | --- | --- |
| 8 | *DEFB43* | 7,449,722 | 7,456,717 |
| 8 | *DEFB134* | 7,469,680 | 7,473,928 |
| 8 | *DEFB136* | 7,499,869 | 7,500,896 |
| 13 | *DEFB132-1* | 60,755,471 | 60,757,652 |
| 13 | *DEFB129-2* | 60,772,966 | 60,775,142 |
| 13 | *DEFB128* | 60,785,898 | 60,787,480 |
| 13 | *DEFB127* | 60,793,742 | 60,796,520 |
| 13 | *DEFB126* | 60,806,515 | 60,811,042 |
| 13 | *DEFB125A* | 60,828,555 | 60,835,043 |
| 13 | *DEFB125* | 60,828,987 | 60,829,208 |
| 13 | *DEFB115* | 60,873,647 | 60,884,399 |
| 13 | *DEFB29* | 60,893,940 | 60,905,009 |
| 13 | *DEFB116* | 60,920,317 | 60,925,563 |
| 13 | *DEFB117* | 60,956,394 | 60,959,265 |
| 13 | *DEFB118* | 60,979,467 | 60,979,700 |
| 13 | *DEFB119-1* | 60,981,163 | 60,990,988 |
| 13 | *DEFB119-2* | 60,989,366 | 60,990,988 |
| 13 | *DEFB121* | 61,007,870 | 61,009,276 |
| 13 | *DEFB122A* | 61,019,572 | 61,023,625 |
| 13 | *DEFB122* | 61,030,365 | 61,034,955 |
| 13 | *DEFB123-2* | 61,038,268 | 61,052,092 |
| 13 | *DEFB123-1* | 61,041,375 | 61,051,806 |
| 13 | *DEFB124* | 61,067,234 | 61,069,998 |
| 23 | *DEFB114* | 22,612,721 | 22,615,376 |
| 23 | *DEFB113* | 22,632,551 | 22,635,067 |
| 23 | *DEFB110-1* | 22,644,099 | 22,656,635 |
| 23 | *DEFB110-2* | 22,651,406 | 22,656,635 |
| 23 | *DEFB112* | 22,662,983 | 22,671,406 |
| 27 | *DEFB1-1* | 5,960,716 | 5,968,525 |
| 27 | *TAP* | 6,013,830 | 6,015,648 |
| 27 | *DEFB103B* | 6,023,993 | 6,025,303 |
| 27 | *SPAG11B* | 6,045,534 | 6,050,941 |
| 27 | *DEFB104* | 6,069,928 | 6,074,858 |
| 27 | *DEFB106A* | 6,076,779 | 6,079,838 |
| 27 | *DEFB15* | 6,076,989 | 6,079,896 |
| 27 | *DEFB105* | 6,082,209 | 6,084,799 |
| 27 | *DEFB107A* | 6,090,910 | 6,095,691 |
| 27 | *DEFB4A* | 7,138,963 | 7,140,882 |
| 27 | *DEFB5* | 7,139,081 | 7,140,848 |
| 27 | *EBD* | 7,165,176 | 7,180,422 |
| 27 | *DEFB130-1* | 6,192,946 | 6,196,744 |
| 27 | *LAP* | 6,234,006 | 6,235,870 |
| 27 | *DEFB33* | 6,238,997 | 6,242,941 |
| 27 | *DEFB13* | 6,326,696 | 6,328,616 |
| 27 | *DEFB103A* | 6,432,334 | 6,433,571 |
| 27 | *DEFB103A-Like* | 6,502,337 | 6,503,278 |
| 27 | *DEFB10* | 6,596,623 | 6,598,299 |
| 27 | *DEFB3* | 6,676,309 | 6,723,692 |
| 27 | *DEFB7* | 6,676,383 | 6,678,273 |
| 27 | *DEFB6* | 6,676,387 | 6,996,057 |
| 27 | *DEFB9* | 6,676,540 | 6,678,161 |
| 27 | *DEFB1-2* | 6,721,772 | 6,723,692 |
| 27 | *DEFB402* | 6,855,327 | 6,856,946 |
| 27 | *DEFB108B* | 7,272,455 | 7,275,948 |
| 27 | *DEFB109* | 7,281,838 | 7,289,252 |

**Supplementary Table 5.**
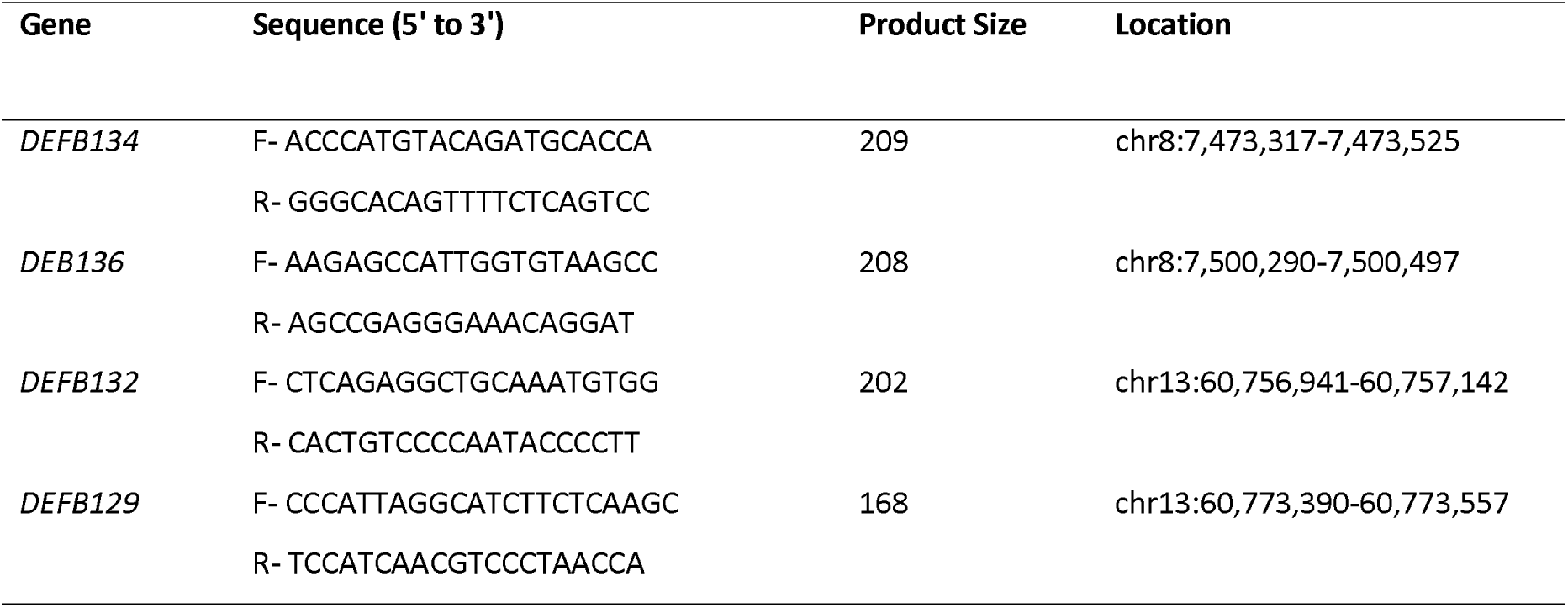
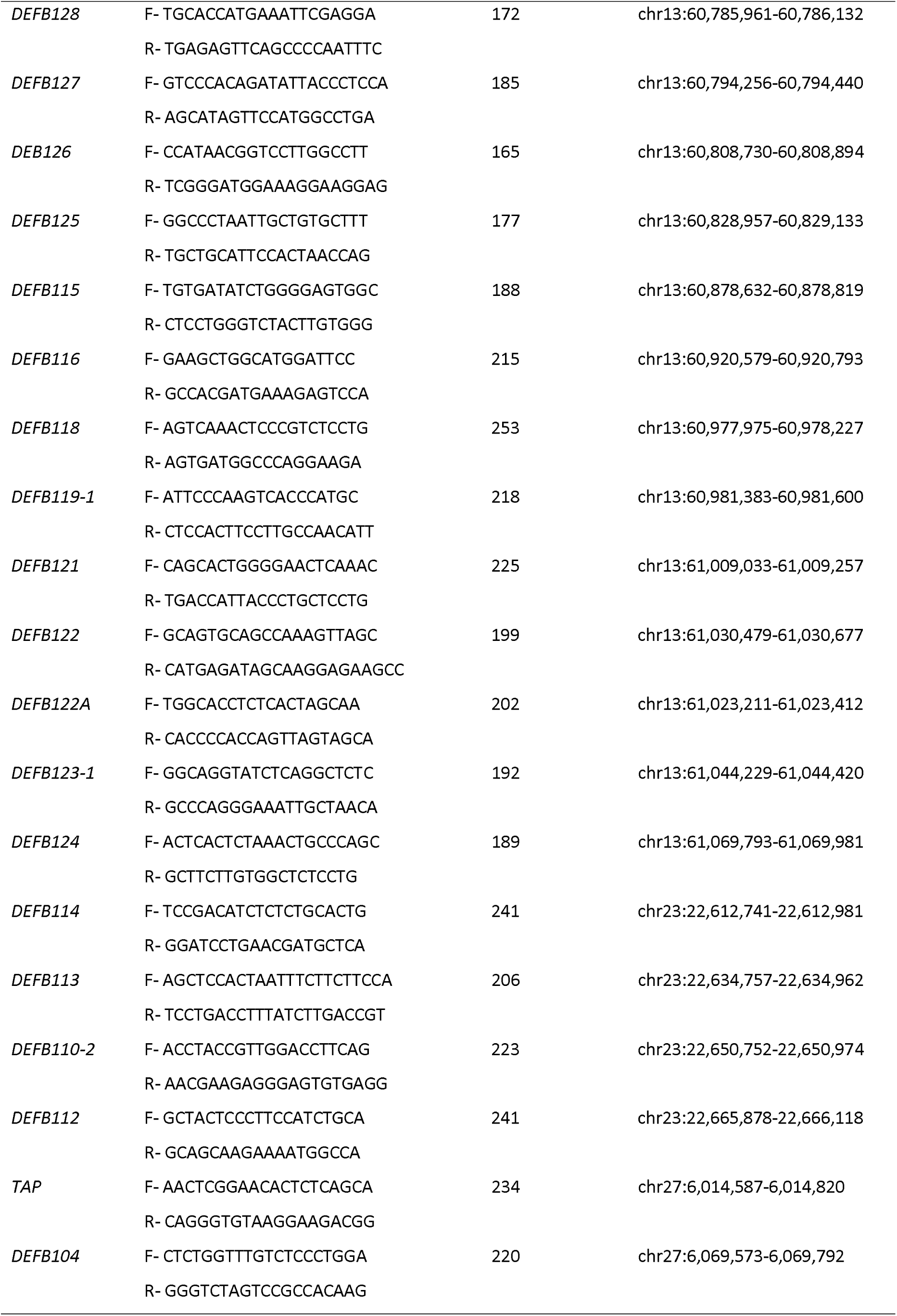
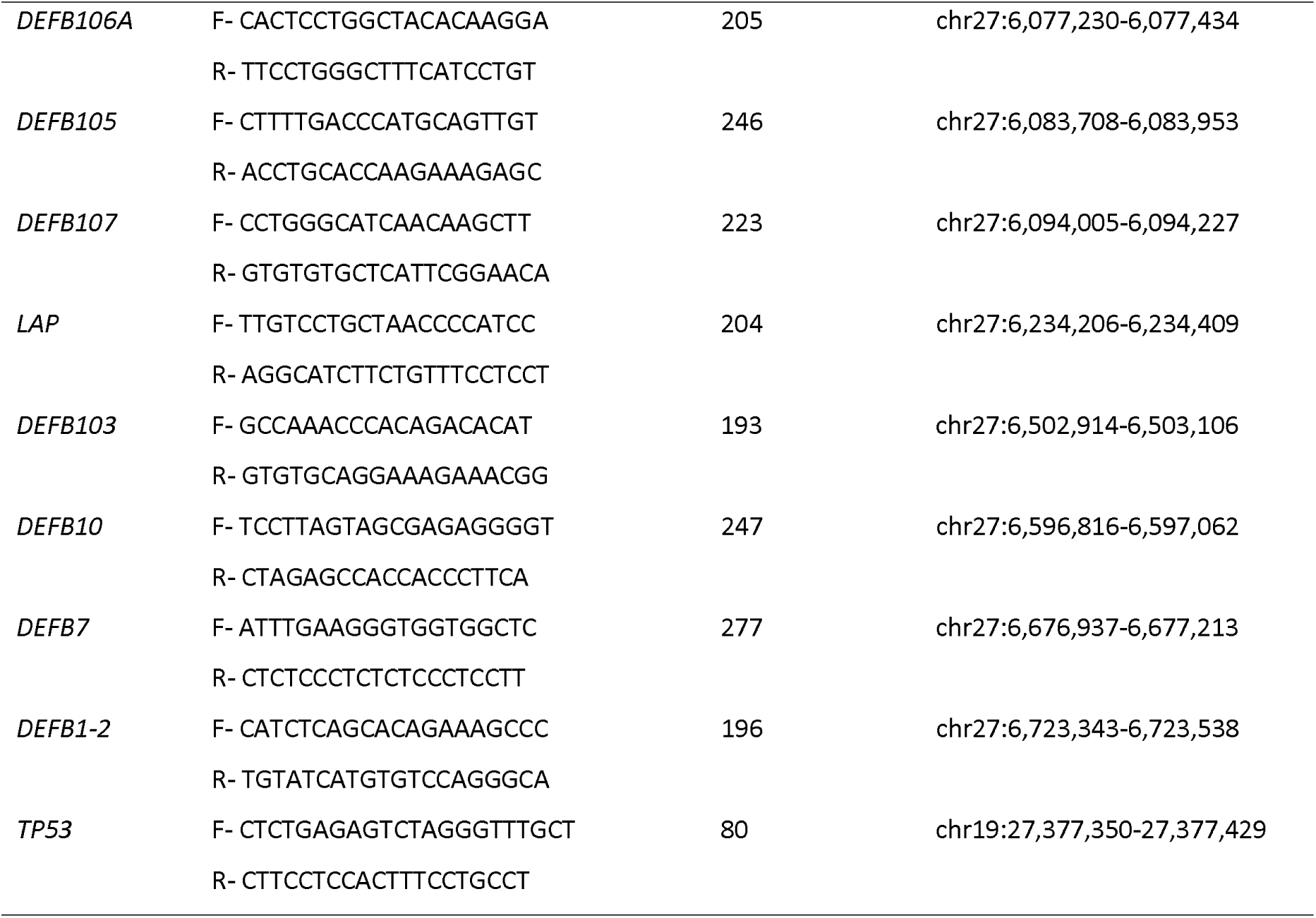
Primer sequences of β-defensin genes analysed using ddPCR.

**Supplementary Table 6.** Β-defensin genes analysed by RNASeq. 46 β-defensins genes were annotated in Ensembl for the bosTau9 genome. The SPAG11 and DEFB genes were excluded.

| Chr | Gene ID | Gene Name | Seq Start | Seq End | Total Size |
| --- | --- | --- | --- | --- | --- |
| 8 | ENSBTAG00000054255 | *DEFB134* | 7469811 | 7472246 | 2436 |
| 8 | ENSBTAG00000051632 | *DEFB136* | 7500173 | 7500855 | 683 |
| 13 | ENSBTAG00000049760 | *DEFB132* | 60755630 | 60757340 | 1711 |
| 13 | ENSBTAG00000048288 | *DEFB129-2* | 60772966 | 60775142 | 2177 |
| 13 | ENSBTAG00000051173 | *DEFB128* | 60785898 | 60787480 | 1583 |
| 13 | ENSBTAG00000054958 | *DEFB127* | 60793877 | 60796412 | 2536 |
| 13 | ENSBTAG00000051033 | *DEFB126* | 60806515 | 60811042 | 4528 |
| 13 | ENSBTAG00000053093 | *DEFB125* | 60824696 | 60832394 | 7699 |
| 13 | ENSBTAG00000049491 | *DEFB115* | 60873655 | 60876483 | 2829 |
| 13 | ENSBTAG00000050556 | *DEFB29* | 60894019 | 60894243 | 225 |
| 13 | ENSBTAG00000052478 | *DEFB116* | 60920397 | 60925535 | 5139 |
| 13 | ENSBTAG00000048009 | *DEFB117* | 60956394 | 60959265 | 2872 |
| 13 | ENSBTAG00000052743 | *DEFB118* | 60972925 | 61008413 | 35489 |
| 13 | ENSBTAG00000003364 | *DEFB119* | 60979366 | 60990988 | 11623 |
| 13 | ENSBTAG00000048949 | *DEFB121* | 61007993 | 61009276 | 1284 |
| 13 | ENSBTAG00000027384 | *DEFB122A* | 61019487 | 61023677 | 4191 |
| 13 | ENSBTAG00000027383 | *DEFB122* | 61030366 | 61034955 | 4590 |
| 13 | ENSBTAG00000020555 | *DEFB123-1* | 61041375 | 61051806 | 10432 |
| 13 | ENSBTAG00000031254 | *DEFB124* | 61067234 | 61069998 | 2765 |
| 23 | ENSBTAG00000054219 | *DEFB114* | 22612852 | 22615351 | 2500 |
| 23 | ENSBTAG00000054396 | *DEFB113* | 22633444 | 22634981 | 1538 |
| 23 | ENSBTAG00000051453 | *DEFB110* | 22651567 | 22656537 | 4971 |
| 23 | ENSBTAG00000046711 | *DEFB112* | 22663626 | 22669740 | 6115 |
| 27 | ENSBTAG00000050611 | *DEFB1-1* | 5960716 | 5968525 | 7810 |
| 27 | ENSBTAG00000048171 | *TAP* | 6013825 | 6015791 | 1967 |
| 27 | ENSBTAG00000052579 | *DEFB103B* | 6023984 | 6025923 | 1940 |
| 27 | ENSBTAG00000034952 | *SPAG11B* | 6063471 | 6068440 | 4970 |
| 27 | ENSBTAG00000048847 | *DEFB104* | 6069928 | 6074858 | 4931 |
| 27 | ENSBTAG00000049070 | *DEFB106A* | 6076779 | 6079838 | 3060 |
| 27 | ENSBTAG00000052911 | *DEFB105A* | 6081510 | 6084079 | 1570 |
| 27 | ENSBTAG00000054607 | *DEFB107A* | 6091050 | 6094942 | 3893 |
| 27 | ENSBTAG00000053557 | *DEFB4A* | 7138873 | 7140876 | 2004 |
| 27 | ENSBTAG00000033545 | *EBD* | 7165176 | 7180420 | 15245 |
| 27 | ENSBTAG00000051223 | *DEFB130* | 6192951 | 6196588 | 3638 |
| 27 | ENSBTAG00000053889 | *LAP* | 6233756 | 6235885 | 2130 |
| 27 | ENSBTAG00000046937 | *DEFB33* | 6238997 | 6242940 | 3944 |
| 27 | ENSBTAG00000050630 | *DEFB13* | 6326521 | 6328629 | 2109 |
| 27 | ENSBTAG00000050419 | *DEFB103A* | 6432444 | 6433381 | 938 |
| 27 | ENSBTAG00000052422 | *DEFB103A-Like* | 6432007 | 6503305 | 71299 |
| 27 | ENSBTAG00000048737 | *DEFB10* | 6596422 | 6598413 | 1992 |
| 27 | ENSBTAG00000051383 | *DEFB7* | 6676076 | 6678281 | 1992 |
| 27 | ENSBTAG00000052312 | *DEFB1-2* | 6721625 | 6723684 | 2060 |
| 27 | ENSBTAG00000053456 | *DEFB402* | 6855327 | 6856946 | 1620 |
| 27 | ENSBTAG00000054169 | *DEFB108B* | 7272744 | 7275810 | 3067 |
| 19 | ENSBTAG00000001069 | *TP53* | 27376072 | 27388416 | 12345 |

**Supplementary Table 7.**
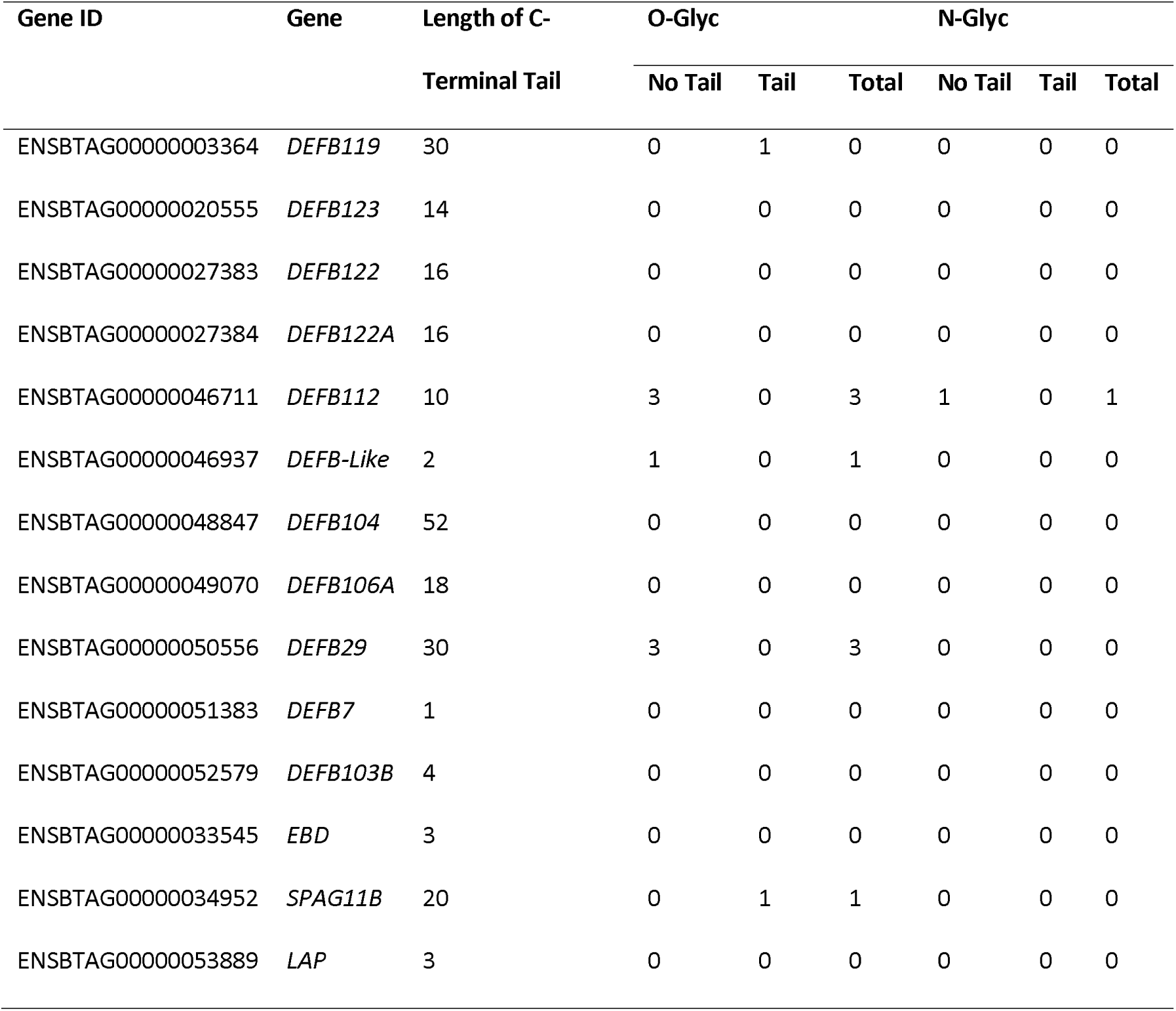
Glycosylation potential and C-terminal tail length of 14 β-defensin proteins upregulated in adult testes.

**Supplementary Figure 1.**
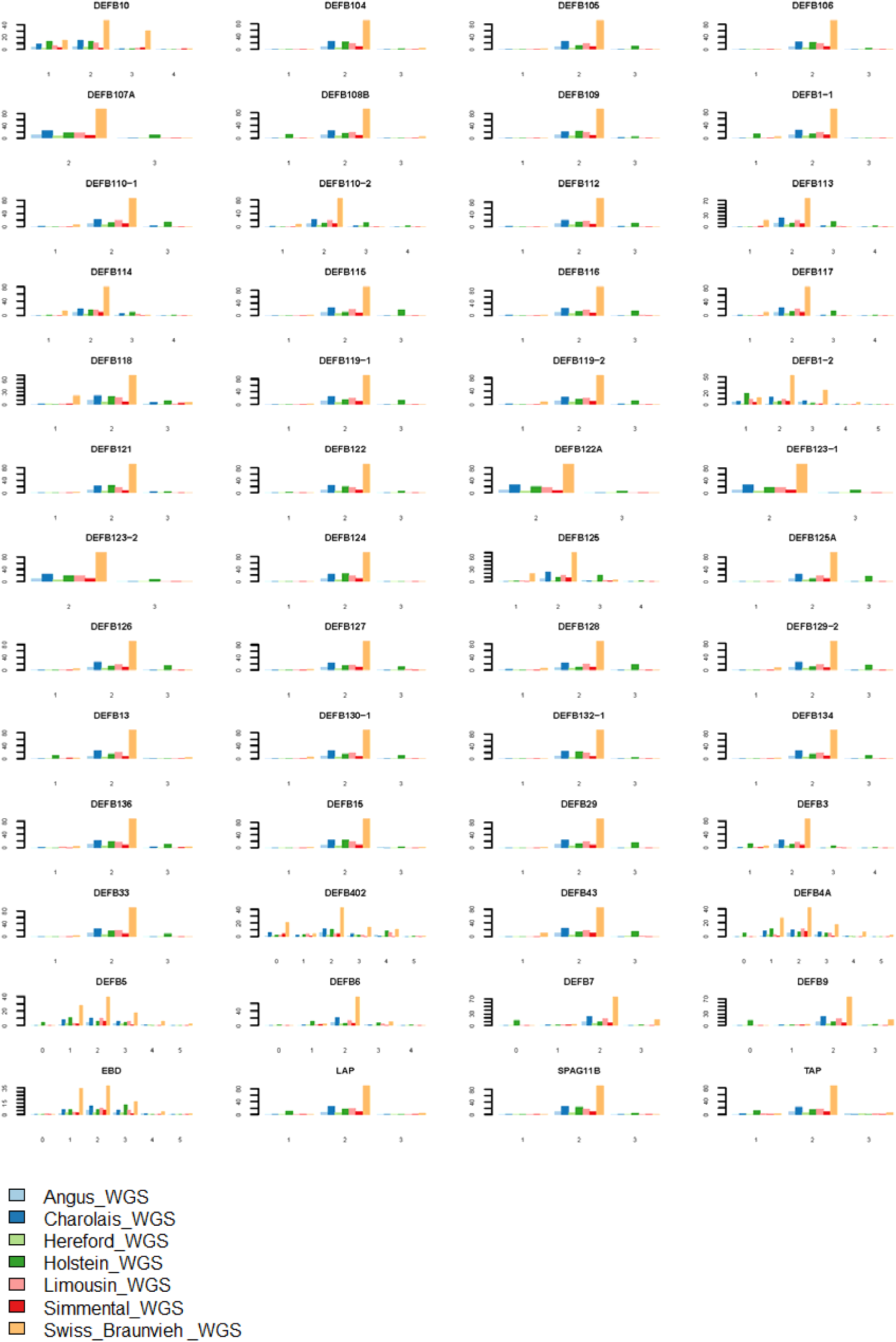
Copy number distribution of different genes.

**Supplementary Figure 2.**
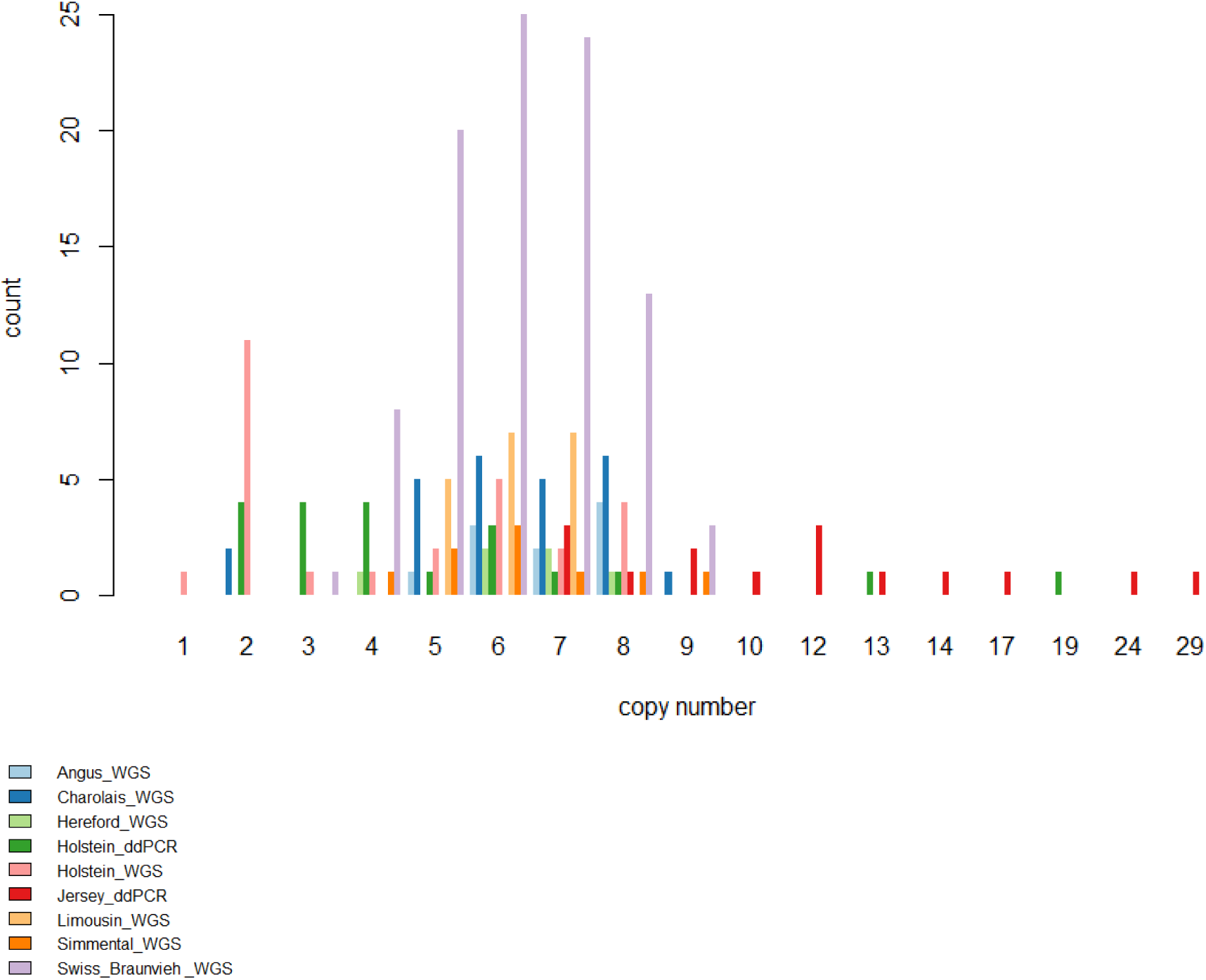
Copy number distribution of DEFB103.

**Supplementary Figure 3.**
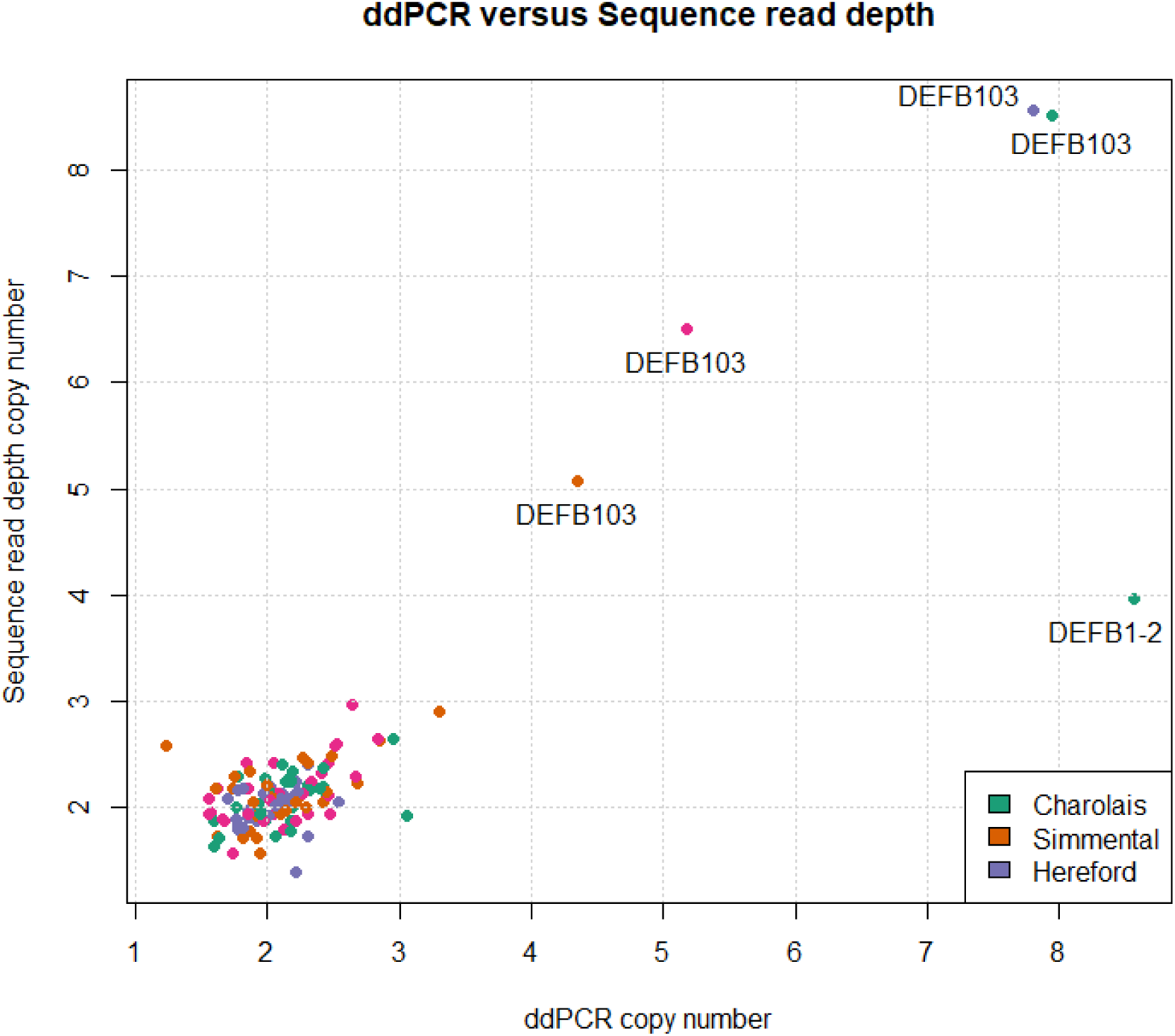
Copy number variation using ddPCR versus WGS read depth.

**Supplementary Figure 4.**
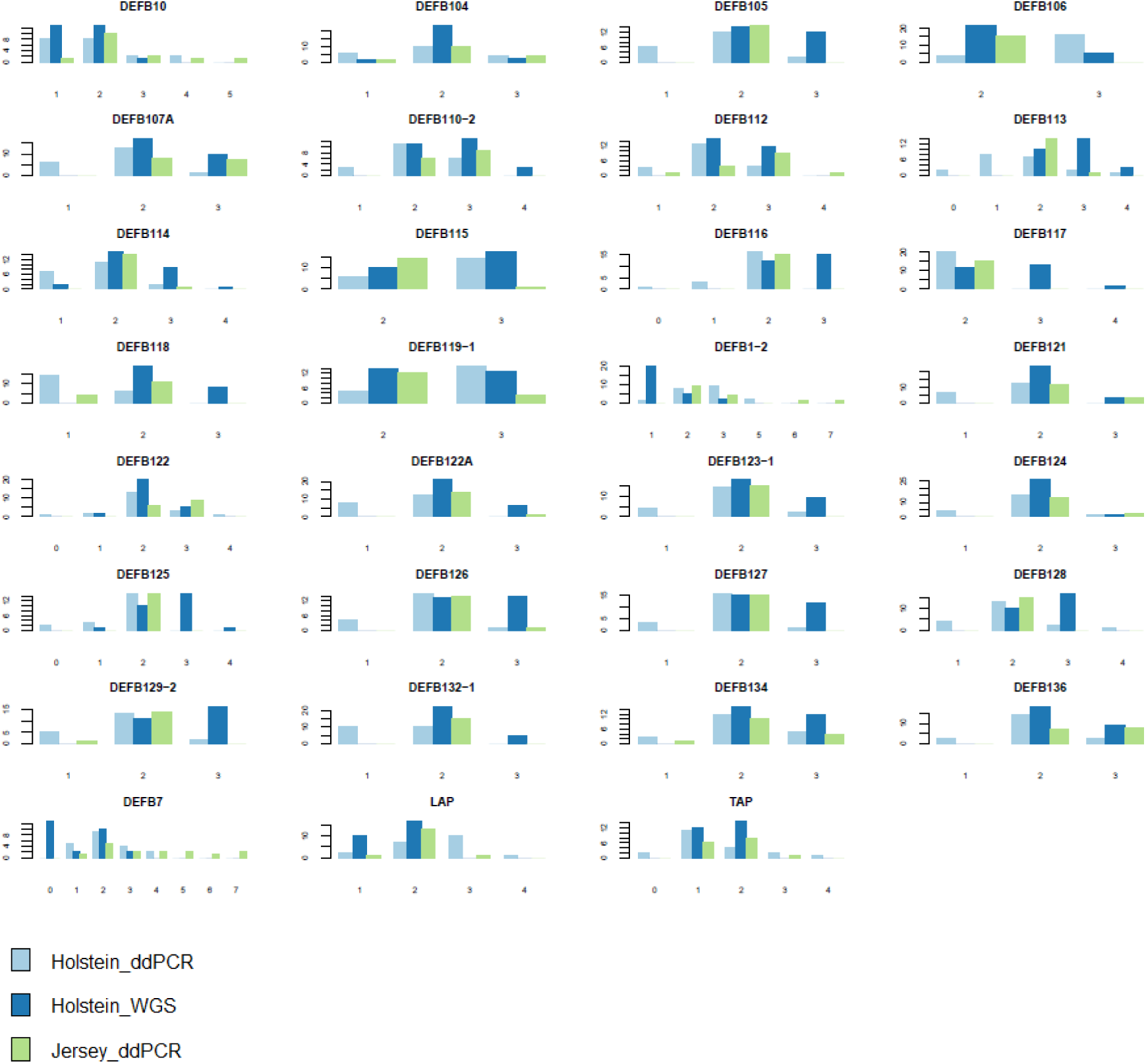
ddPCR analysis of copy number for Holstein and Jersey.

## Notes

https://doi.org/10.25392/leicester.data.25992484.v1

https://doi.org/10.25392/leicester.data.25991863.v1

### Competing Interest Statement

The authors have declared no competing interest.

